# Multi-omic analysis of the Arabidopsis clock activator mutant *rve 4 6 8* reveals connections to carbohydrate metabolism and proteasome regulation

**DOI:** 10.1101/2021.10.25.465654

**Authors:** S Scandola, D Mehta, Q Li, M Rodriguez, B Castillo, RG Uhrig

**Author notes:** Corresponding Author, R. Glen Uhrig, Department of Biological Sciences, University of Alberta, 11455 Saskatchewan Drive, Edmonton, Alberta, Canada, T6G 2E9. denotes equal authorship (S Scandola, D Mehta).

## Abstract

Plants are able to sense changes in their light environments, such as the onset of day and night, as well as anticipate these changes in order to adapt and survive. Central to this ability is the plant circadian clock, a molecular circuit that precisely orchestrates plant cell processes over the course of a day. REVEILLE proteins (RVEs) are recently discovered members of the plant circadian circuitry that activate the evening complex and PRR genes to maintain regular circadian oscillation. The RVE 8 protein and its two homologs, RVE 4 and 6, have been shown to limit the length of the circadian period, with *rve 4 6 8* triple-knockout plants possessing an elongated period along with increased leaf surface area, biomass, cell size and delayed flowering relative to wild-type Col-0 plants. Here, using a multi-omics approach consisting of phenomics, transcriptomics, proteomics, and metabolomics we draw novel connections between RVE8-like proteins and a number of core plant cell processes. In particular, we reveal that loss of RVE8-like proteins results in altered carbohydrate, organic acid and lipid metabolism, including a starch excess phenotype at dawn. We further demonstrate that *rve 4 6 8* plants have lower levels of 20S proteasome subunits and possess significantly reduced proteasome activity, potentially explaining the increase in cell-size observed in RVE8-like mutants. Overall, this robust, multi-omic dataset, provides substantial new insights into the far reaching impact RVE8-like proteins have on the diel plant cell environment.

## INTRODUCTION

The circadian clock is a central regulator of plant growth and development that modulates plant responses to both internal and external cues (Greenham and McClung, 2015; Nohales and Kay, 2016; McClung, 2019). It is comprised of a number of transcription factors that function as a series of interlocking molecular circuits that precisely time the 24-hour photoperiod (Creux and Harmer, 2019). Correspondingly, knockout mutants of circadian clock transcription factors have been extensively studied at the phenotypic, genetic and transcript-level in the model plant *Arabidopsis thaliana* (Creux and Harmer, 2019; McClung, 2019; Nakamichi, 2020), with upwards of 30% of genes shown to be under circadian control (Covington et al., 2008). This extensive body of work has revealed a number of roles for the circadian clock in plants, including in the timing of flowering (Shim et al., 2017), disease resistance (Lu et al., 2017) and mitigation of abiotic stress (Simon et al., 2020; Kidokoro et al., 2021) amongst others. However, despite the extensive use of genetic and transcriptomic technologies to understand the molecular impacts of core circadian clock components in *Arabidopsis*, much less is known about how, and/or if, these transcriptional changes manifest at the protein-level (Choudhary et al., 2015; Choudhary et al., 2016; Krahmer et al., 2022; Uhrig et al., 2019; Mehta et al., 2021; Uhrig et al., 2021).

Protein abundance and post-translational protein modification (PTM) changes are critical control mechanisms for biological systems. To date, quantitative proteomic analysis of circadian clock mutants *lhy cca1*, *prr7 prr9*, *gi* and *toc1* at end-of-day (ED; ZT12) and end-of-night (EN; ZT0) time-points (Graf et al., 2017) and time-course experimentation involving CCA1_ox_ (Krahmer et al., 2022) and wild-type (WT) Col-0 (Uhrig et al., 2021) plants over free-running and 12:12 light:dark (LD) conditions have provided the first insights into where diel changes in the proteome manifest. Similarly, time-course and ED versus EN analyses of the phosphoproteome from CCA1_ox_ (Krahmer et al., 2022) and WT Col-0 (Uhrig et al., 2019) plants point to a critical, but largely unexplored, role for protein phosphorylation and other PTMs in diel plant cell regulation; particularly at ED and EN. Beyond proteomics, quantitative metabolomic analyses of core circadian clock mutants can also help define the role the circadian clock plays in regulating growth and development through the modulation of primary metabolism (Fukushima et al., 2009; Flis et al., 2019).

REVEILLE (RVE) genes are recently discovered additions to the circadian clock that have been found to function as core circadian clock activators, modulating the regulation of the evening complex (EARLY FLOWERING 3 (ELF3), 4 (ELF4) and LUX ARRHYTHMO (LUX)) in addition to TIMING OF CAB EXPRESSION 1 (TOC1) and PSEUDO RESPONSE REGULATOR 5 (PRR5) (Farinas and Mas, 2011; Rawat et al., 2011; Hsu et al., 2013; Gray et al., 2017). Findings that *rve 8* (Farinas and Mas, 2011; Rawat et al., 2011) and *rve 4 6 8* (Gray et al., 2017) plants possess an elongated circadian period demonstrate how RVE proteins are important to maintaining the pace of the circadian clock, while phenotypic characterization of the *rve 4 6 8* plants revealed greater plant growth and cell size, coupled with a delay in flowering (Gray et al., 2017).

Despite these findings, how RVE8-like proteins regulate the diel plant cell environment at the protein- and metabolic-level remains unresolved. To address this gap in knowledge, we have undertaken a multi-omic analysis of *rve 4 6 8* plants relative to WT Col-0 at critical diel / circadian time-points; EN / ZT0 and ED / ZT12. Here, we demonstrate that loss of RVE 4, 6 and 8 results in extensive diel proteome, phosphoproteome and metabolome changes spanning a wide array of critical plant cell processes. In particular, our results suggest that RVE8-like proteins impact critical elements of primary metabolism such as transient diel leaf-starch / carbohydrate levels. Perhaps most intriguingly, we find that *rve 4 6 8* plants maintain extensive protein- and PTM-level perturbations in proteasome subunits, coupled with reduced proteasome activity and altered amino acids levels, which together, likely form the basis of the increased growth phenotype of *rve 4 6 8*. As RVE8-like proteins are associated with multiple agronomically important traits such as biomass and flowering time, elucidating their role in modulating the diel proteome, PTMome and metabolome is critical for their potential implementation in crop improvement strategies.

## RESULTS

### Examination of *rve 4 6 8* mutant phenotypes reveals starch excess at dawn

In order to ensure our molecular analysis would provide robust results, we first phenotypically characterized *rve 4 6 8* plants relative to WT Col-0 (Col-0) under our growth conditions (12:12 LD photoperiod facilitated by LED lights). Here, we were able to reproduce multiple growth and development phenotypes previously observed (Gray et al., 2017), indicating that our growth conditions are comparable (Supplemental Figure 1). We monitored plant area and perimeter using real-time imaging over 5 days and found significant differences between *rve 4 6 8* and WT Col-0 arise by day 17 post-imbibition, with *rve 4 6 8* plants possessing increased leaf area and perimeter (Supplemental Figure 1A & B). We then performed fresh and dry weight measurements of *rve 4 6 8* versus WT Col-0 plants at day 19 post-imbibition revealing that *rve 4 6 8* has greater biomass (Supplemental Figure 1C & D). Lastly, under our 12:12 LD conditions, we found *rve 4 6 8* plants have delayed flowering relative to WT Col-0 (Supplemental Figure 1E).

Given these growth phenotypes and the previously observed response of *rve 4 6 8* to sucrose (Gray et al., 2017), we next stained *rve 4 6 8* and WT Col-0 rosettes for the accumulation of transient leaf starch over a 12:12 LD photoperiod examining ZT0, 6, 12 and 18 time-points. This revealed that *rve 4 6 8* plants possess a starch excess phenotype at ZT0 (Figure 1A). We then validated this analysis by performing an enzymatic quantitation of starch levels in WT Col-0 and *rve 4 6 8* plants (Figure 1B). Samples were collected hourly from ZT21 to ZT3 in order to enzymatically assess if the starch excess observed was due to the long-period phase shift of the *rve 4 6 8* mutant clock. Here, our results indicate that the observed starch excess is in fact independent of the long-period phase shift of the *rve 4 6 8* mutant (Figure 1B). We then further examined three key starch metabolism related enzymes for changes in their gene expression using NanoString RNA quantification: *PHOSPHOGLYCERATE MUTASE 1* (*PGM1*), *GRANUAL BOUND STARCH SYNTHASE 1* (*GBSS1*) and *STARCH EXCESS 4* (*SEX4*). *GBSS1* and *SEX4* transcript levels were significantly lower in *rve 4 6 8* compared to WT Col-0 at ZT0 and ZT12, respectively, while no change was observed in PGM expression (Figure 1C and D). Together with our phenotypic data, these results suggest a potential disconnect between the timing of starch degradation and biosynthesis that results in greater transient starch accumulation at dawn.

**Figure 1:**
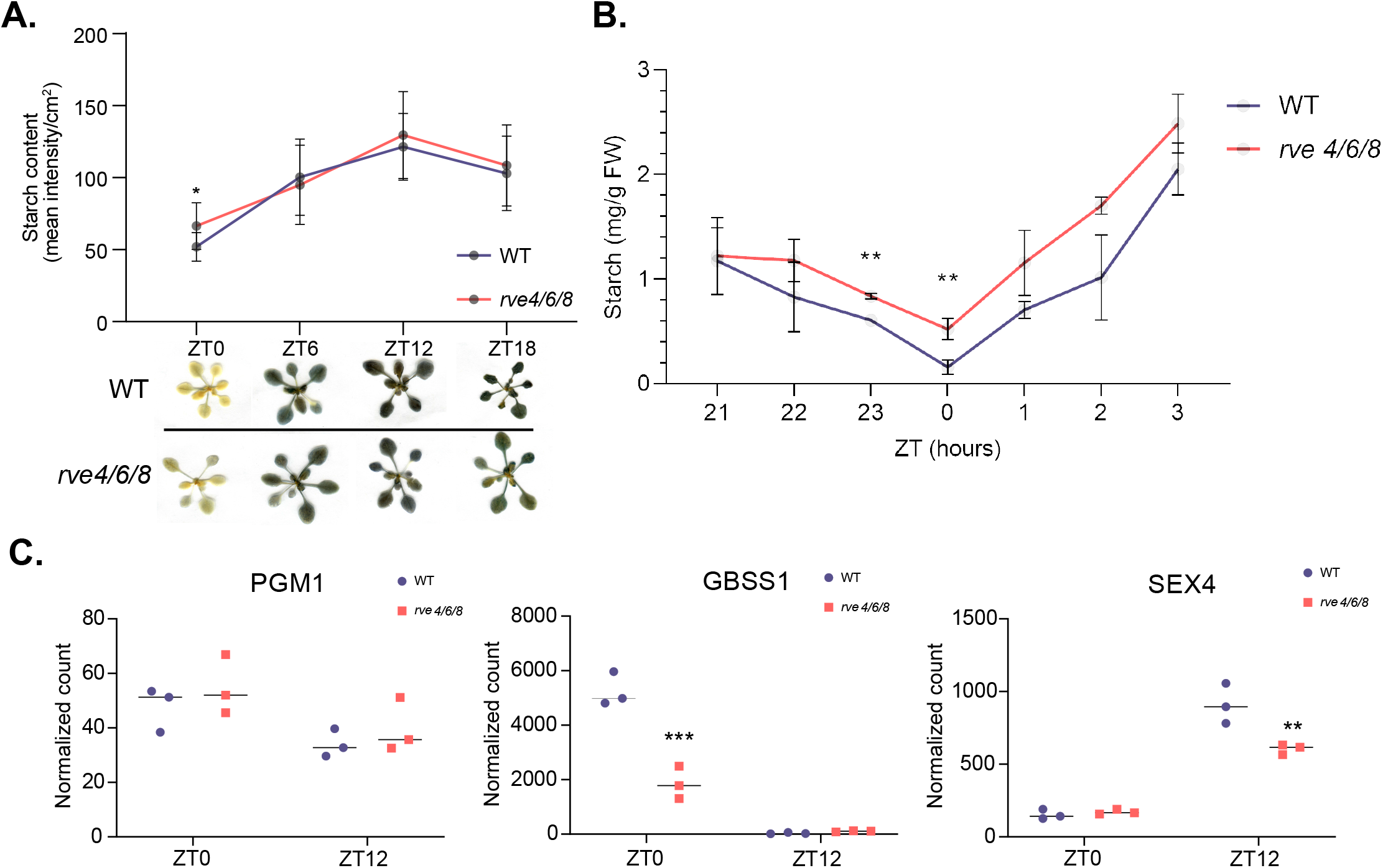
Phenotypic analysis of *rve 4 6 8* plant growth and development. **(A)** Iodine starch staining of rosettes at 28 d post-imbibition. (**B**) Enzymatic quantification of starch levels in rosettes. All phenotyping was conducted under a 12 h : 12 h light : dark photoperiod using 100 μmol m-2 s-1 light. Stars denote Bonferroni adjusted *p-value* significance: (***) *p-value* ≤ 0.005 and (*) *p-value* ≤ 0.05. **(C)** Starch related genes PGM1, GBSS1 and SEX4. All time-points and genotypes were analyzed in biological triplicate (n=3). Stars denote Student’s t-test *p-value* significance: (***) *p-value* ≤ 0.005, (**) *p-value* ≤ 0.01 and (*) *p-value* ≤ 0.05. Primer pairs used in the nanostring analysis are described in Supplemental Data 1.

### Gene expression analysis provides molecular insights into *rve 4 6 8* growth phenotypes

Using NanoString RNA quantification, we next examined the expression of a panel of core circadian clock genes and genes involved in processes impacted by the circadian clock (Figure 2; Supplemental Figure 2 Supplemental Data 1). Studies examining RVE 8-like proteins (RVE 4, 6 & 8) have observed changes in multiple core circadian clock and clock-associated genes (Creux and Harmer, 2019), including *PSEUDO RESPONSE REGULATOR 5* (*PRR5*), *TIMING OF CAB1 EXPRESSION 1* (*TOC1*), *EARLY FLOWERING 4* (*ELF4*), *PHYTOCHROME INTERACTING FACTOR 4* (*PIF4*) and *5* (*PIF5*). We aimed to consolidate this understanding by examining changes in these genes, amongst others, in *rve 4 6 8* plants at ZT0 and ZT12 time-points under 12:12 LD conditions. Here, we find a decrease in *TOC1* and *ELF4* transcript levels combined with an increase in *PRR5*, *PIF4* and *PIF5* expression in *rve 4 6 8* at ZT12 (Figure 2A). We also observed a decrease in *CIRCADIAN CLOCK REGULATED 1* (*CCA1*) and *LATE ELONGATED HYPOCOTYL* (*LHY*) expression at ZT0 along with increases in *PSEUDO RESPONSE REGULATOR 7* (*PRR7*) and *GIGANTEA* (*GI*) at ZT12 that are also consistent with an elongated period (Hsu et al., 2013; Flis et al., 2016). While our PRR5 result accords with previous findings, our Nanostring quantification shows reduced TOC1 transcript levels while previous qPCR results have shown increased TOC1 levels at ZT12 (Hsu et al, 2013). Similarly, while we observe reduced transcript levels of LHY and CCA1 in the mutant, the previous study does not observe any difference. These differences between the two studies is likely due to the different measurement techniques used (qPCR vs Nanostring) and the relatively larger variation in expression recorded in Hsu et al. 2013. Collectively, our results align with previous research examining RVE8-like proteins, however, more time-points are needed to precisely define the impact of RVE8-like proteins on the phasing and period lengths of core clock transcripts.

**Figure 2:**
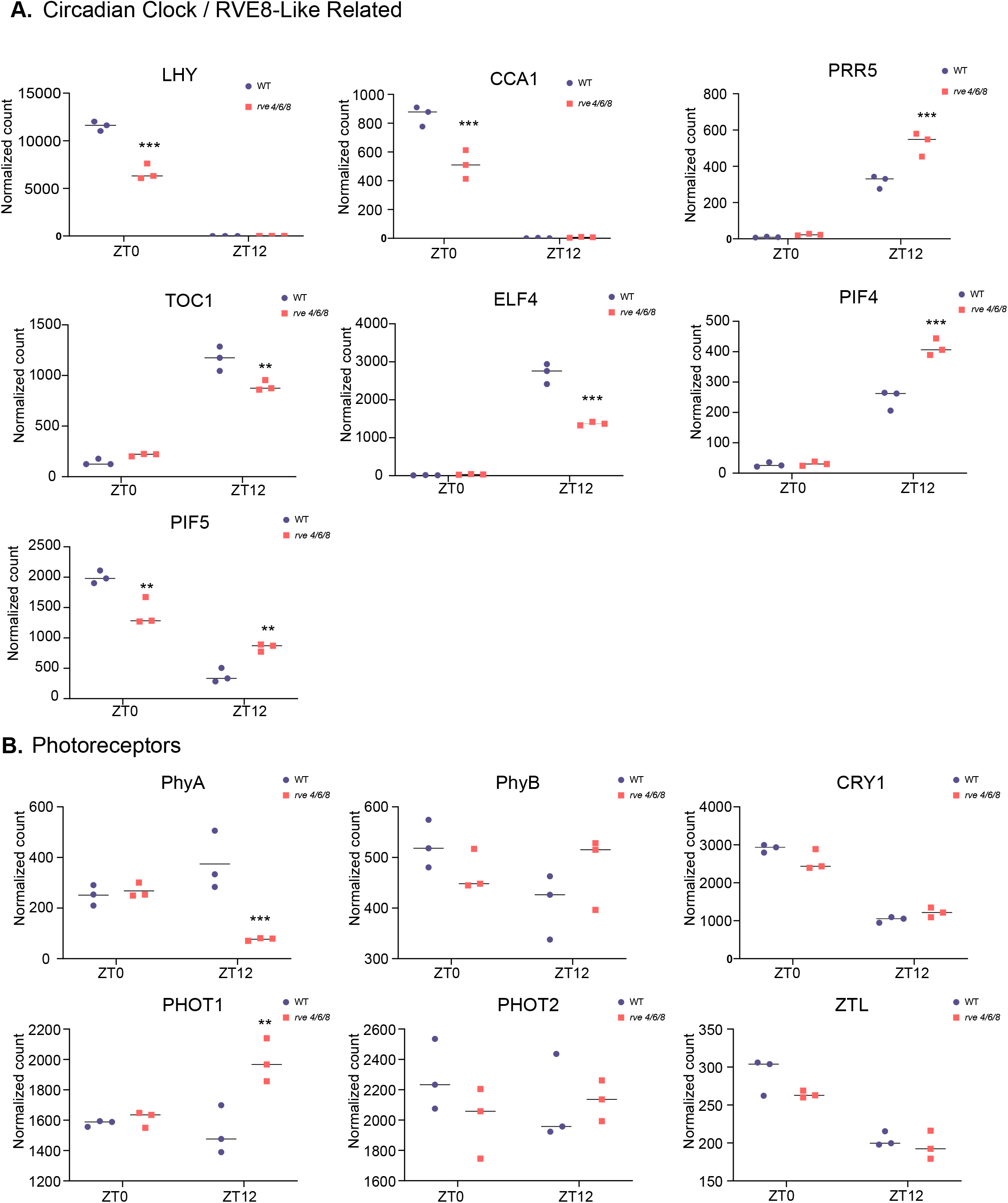
Diel mRNA gene expression analysis of circadian clock and photoreceptor related genes in *rve 4 6 8* and Col-0 plants. Select genes from different circadian / diel implicated plant cell processes were examined for their expression level at end-of-night (ZT0) and end-of-day (ZT12) by NanoString expression analysis (see Materials and Methods). **(A)** Expression of circadian clock / RVE8-like related genes *CCA1*, *LHY*, *TOC1*, *PRR5*, *ELF4*, *PIF4* and *PIF5*. **(B)** Expression of photoreceptors *PhyA*, *PhyB*, *CRY1*, *PHOT1*, *PHOT2* and *ZTL*. All time-points and genotypes were analyzed in biological triplicate (n=3). Stars denote Student’s t-test *p-value* significance: (***) *p-value* ≤ 0.005, (**) *p-value* ≤ 0.01 and (*) *p-value* ≤ 0.05. Primer pairs used in the nanostring analysis are described in Supplemental Data 1.

Photoreceptors are crucial in circadian clock entrainment by light, but are in turn also controlled by the clock (Bognar et al., 1999; Sanchez et al., 2020). Correspondingly, we next examined *rve 4 6 8* plants for changes in the expression of key photoreceptor genes, quantifying changes in the expression of *PHYTOCHROME* (*PHY*) *A* and *B*, *PHOTOTROPIN* (*PHOT*) *1* and *2*, *CRYPTOCHROME 1* (*CRY1*) and the circadian clock associated photoreceptor *ZEITLUPE* (*ZTL*) in *rve 4 6 8* versus WT Col-0 at ZT0 and ZT12. Here, we found a significant decrease in *PHYA* and a significant increase in *PHOT1* expression in *rve 4 6 8* plants at ZT12, with no change in *PHYB*, *PHOT2*, *CRY1* and *ZTL* (Figure 2B).

Since *rve 4 6 8* plants demonstrate a number of growth phenotypes, we additionally examined the expression of genes upstream of, or involved in, phytohormone related-processes. Here, we find significant changes in *GERANYLGERANYL PYROPHOSPHATE SYNTHASE 1* (*GGPPS1*) at both ZT0 and ZT12 as well *ARABIDOPSIS RESPONSE REGULATOR 7* (*ARR7*) at ZT12 (Supplemental Figure 2). While these genes represent only a sampling of the phytohormone metabolism / signaling landscape, both their enzyme products have a common connection to Dimethylallyl Pyrophosphate (DMAPP) / Isoprenyl Pyrophosphate (IPP), a critical metabolic intermediate. GGPPS1 catalyzes the entry point reaction for downstream production of multiple phytohormones, including: abscisic acid, gibberellin and stringolactone biosynthesis (Ruiz-Sola et al., 2016). Cytokinin is also derived from the DMAPP / IPP metabolic branch point (Astot et al., 2000) and ARR7 represents a cytokinin response factor that is activated in the presence of cytokinins (Huang et al., 2017).

### Quantitative proteome and phosphoproteome analyses reveal extensive protein-level impacts of RVE8-like protein loss

Quantifying proteome-level changes in the plant cell environment in response to perturbation (e.g. gene deletion) is a robust means of measuring the impact of a protein on a system. However, the high dynamic range nature of plant tissues has typically hampered our ability to sufficiently sample the dynamically fluctuating proteome, biasing analyses towards high abundant proteins (Mehta et al., 2022; Mehta et al., 2021). Given this, we deployed a combination of BoxCarDIA (proteome abundance) and conventional DDA (phosphoproteomics) mass spectrometry acquisition methods to perform label-free quantitation (LFQ) of protein abundance and phosphoproteomic changes in *rve 4 6 8* plants at ZT0 and ZT12. The ZT0 and ZT12 time-points were selected for analysis as they have been shown to exhibit dynamic protein-level changes (e.g. abundance and PTMs) across multiple plant organs, tissues and growth conditions in a variety of wild-type and circadian clock mutant plants (Graf et al., 2017; Krahmer et al., 2022; Uhrig et al., 2019; Uhrig et al., 2021). Through this analysis we quantified 3,934 proteins and 1,137 phosphoproteins for changes in abundance and phosphorylation status, of which 536 and 346 demonstrated significant fluctuations, respectively (Table I; Supplemental Data 2 and 3). Further, we quantified 46% of all proteins exhibiting a significant change in their phosphorylation status at the proteome-level (159 / 347 phosphoproteins), with 8.6% of phosphoproteins demonstrating a corresponding change in protein abundance (30 / 347 phosphoproteins).

**Table I:** Quantified and significantly changing proteome and phosphoproteome. Protein groups and peptides are shown (n=4; *q-value* ≤ 0.05; fold-change (FC) ≥ 1.5).

|  | Quantified |  | *Significantly Changing |  |
| --- | --- | --- | --- | --- |
|  | Protein Groups | Peptides | Protein Groups | Peptides |
| Proteome | 3934 | n/a | 536 | n/a |
| Phosphoproteome | 1137 | 1872 | 346 | 425 |

\* (n=4; *q-value* ≤ 0.05; FC ≥ 1.5)

Next, using significantly changing proteins and phosphoproteins, we performed a gene ontology (GO) enrichment analysis for biological processes, subcellular localization and molecular function to reveal how the loss of RVE8-like proteins impacts the diel plant cell environment at the protein-level (Figure 3; Supplemental Data 4). Here, we found changes in ‘proteolysis’ (GO:0006508), which was complemented by the enrichment of ‘proteasome core complex’ (GO:0005839) and ‘proteasome complex’ (GO:0000502) subcellular localizations, in addition to ‘response to acid chemical’ (GO:0001101), which represents a high-level category for phytohormones and ‘polysaccharide metabolism’ (GO:0005976), amongst others (Figure 3A). Similarly, phosphoproteomic data revealed enrichment in ‘response to acid chemical’ (GO:0001101) and ‘response to abscisic acid’ (GO:0009737) in addition to ‘cell cycle’ (GO:0007049), ‘cell division’ (GO:0051301), ‘macromolecule metabolism’ (GO:0043170) and ‘response to lipids (GO:0033993), amongst others (Figure 3B). While proteome-level *rve 4 6 8*-dependent changes included both increases and decreases in protein levels, we found a global increase in protein phosphorylation in the *rve 4 6 8* mutant compared to WT Col-0 (Figure 3B).

**Figure 3:**
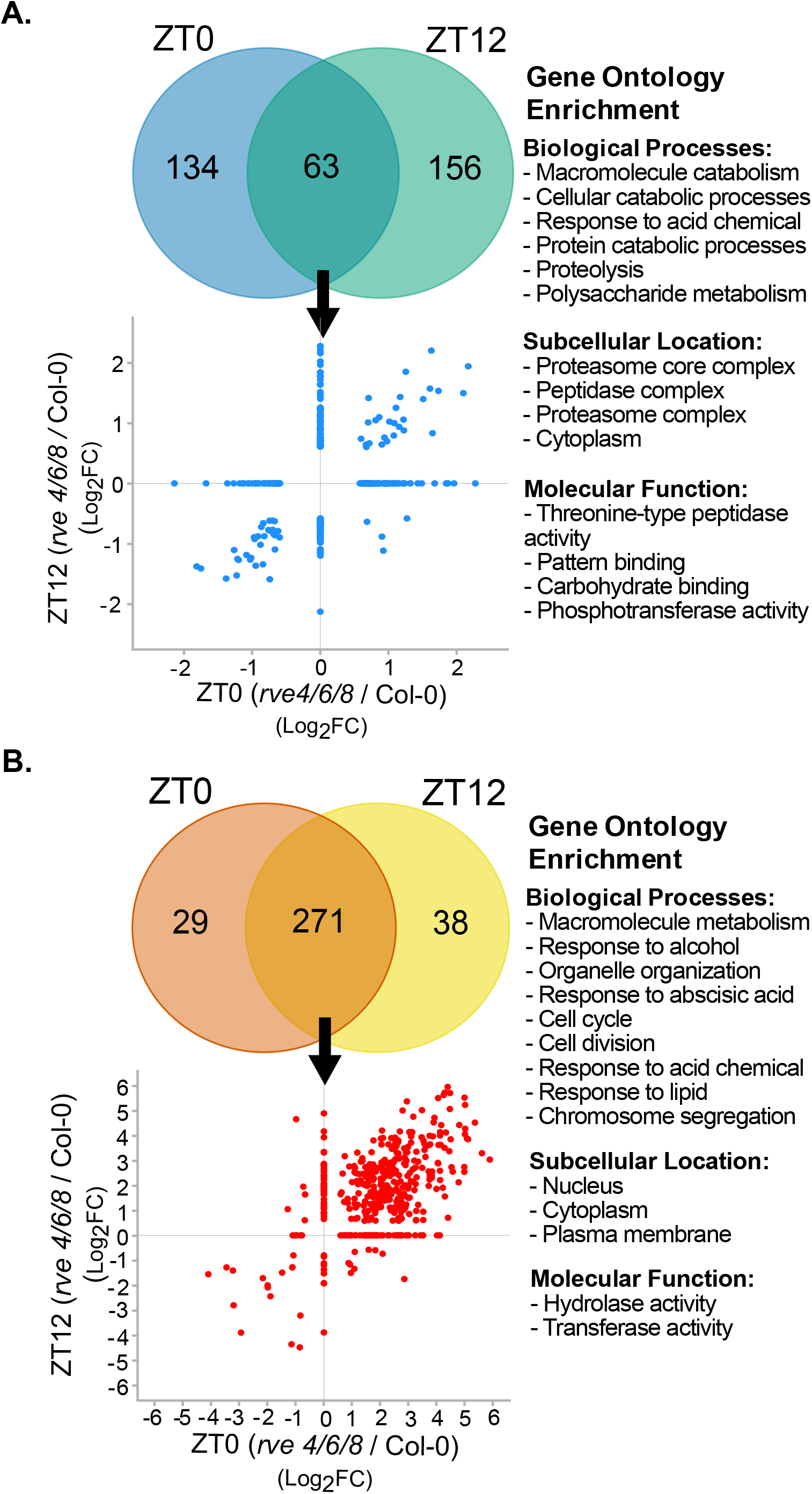
Analysis of the quantified proteome and phosphoproteome changes at ZT0 or ZT12 in *rve 4 6 8* versus Col-0. The significantly changing **(A)** proteome and **(B)** phosphoproteome (n=4; fold-change ≥ 1.5, *q-value* ≤ 0.05) were plotted by venn diagram and scatterplot to visualize the overlap and Log_2_ fold-change (Log_2_FC) between *rve 4 6 8* and WT Col-0 plants at ZT0 and ZT12. Gene ontology (GO) enrichment of biological processes, subcellular localization and molecular function was then performed to systemically contextualize proteome and phosphoproteme changes (see Materials and Methods).

Lastly, we compared our significantly changing phosphopeptides at ZT0 and ZT12 with phospho-sites previously found to oscillate under free-running conditions in WT Col-0 (Krahmer et al., 2022). Here we found that 77 of our phosphopeptides (*q-value* ≤ 0.05) significantly changed in *rve 4 6 8* compared to WT Col-0 (JTK *p-value* ≤ 0.05; Supplemental Data 5). These oscillating phosphopeptides are from diverse, but critically important proteins such as nuclear localized RETINOBLASTOMA-RELATED 1 (RBR; AT3G12280) – growth and development, cytosolic PHOSPHOENOLPYRUVATE CARBOXYLASE 2 (PEPC2; AT2G42600) and chloroplast-targeted RUBISCO ACTIVASE (AT2G39730) - metabolism and plasma membrane associated CELLULOSE SYNTHASE-LIKE D3 (CSLD3; AT3G03050) – cell wall biosynthesis (Supplemental Data 5). This indicates that RVE8-like proteins influence the timing of diel phosphorylation events on proteins involved in core growth and development processes.

### Time-of-day association network analysis finds multiple gene networks affected in *rve 4 6 8* at the protein-level

To contextualize the changing proteome and phosphoproteome, we next generated STRING-DB association networks (minimum edge score ≥ 0.7) for all significantly changing proteins and phosphoproteins in *rve 4 6 8* plants at ZT0 and ZT12 relative to WT Col-0 to better understand the protein-level changes that underpin observed *rve 4 6 8* phenotypes. Proteome quantification revealed decreases in the abundance of multiple starch degradation enzymes including STARCH EXCESS 1 (SEX1; AT1G10760) and SEX4 as well as cold-stress proteins COLD-REGULATED 15A (COR15A; AT2G42540), 15B (COR15B; AT2G42530) and 6 (COR6; AT5G15970). Conversely, we observed increases in primary metabolic and sulfur assimilation enzymes such as PHOSPHOFRUCTOKINASE 5 (PFK5; AT2G22480), PYRUVATE ORHOPHOSPHATE DIKINASE (PPDK; AT4G15530) 5’-ADENYLYLPHOSPHOSULFATE REDUCTASE 2 (APR2; AT1G62180) and ADENYLYL-SULFATE KINASE (APK; AT2G14750), along with a multitude of spliceosomal, ribosomal and ubiquitin ligating proteins. Together, these proteomic data support our observed starch excess phenotype in *rve 4 6 8* at ZT0, while suggesting RVE8-like proteins also impact agronomically important processes such as sulfur assimilation and cold stress response (Figure 4). We also observed distinct changes in proteasome complex proteins, whereby four 19S proteasome proteins (AT1G04810, AT1G75990; AT4G19006, AT4G24820) are up-regulated and twelve 20S proteasome proteins are down-regulated in *rve 4 6 8* plants (Figure 4; Supplemental Data 6). Further examination of the transcripts corresponding to the proteasome subunits changing at the protein-level using DiurnalDB (http://diurnal.mocklerlab.org/) found that 9 of the 12 20S subunits and 2 of the 4 19S subunits possess daily oscillations in their transcripts (≥ 0.8 cutoff), with most peaking in abundance at ZT12 (Supplemental Data 6).

**Figure 4:**
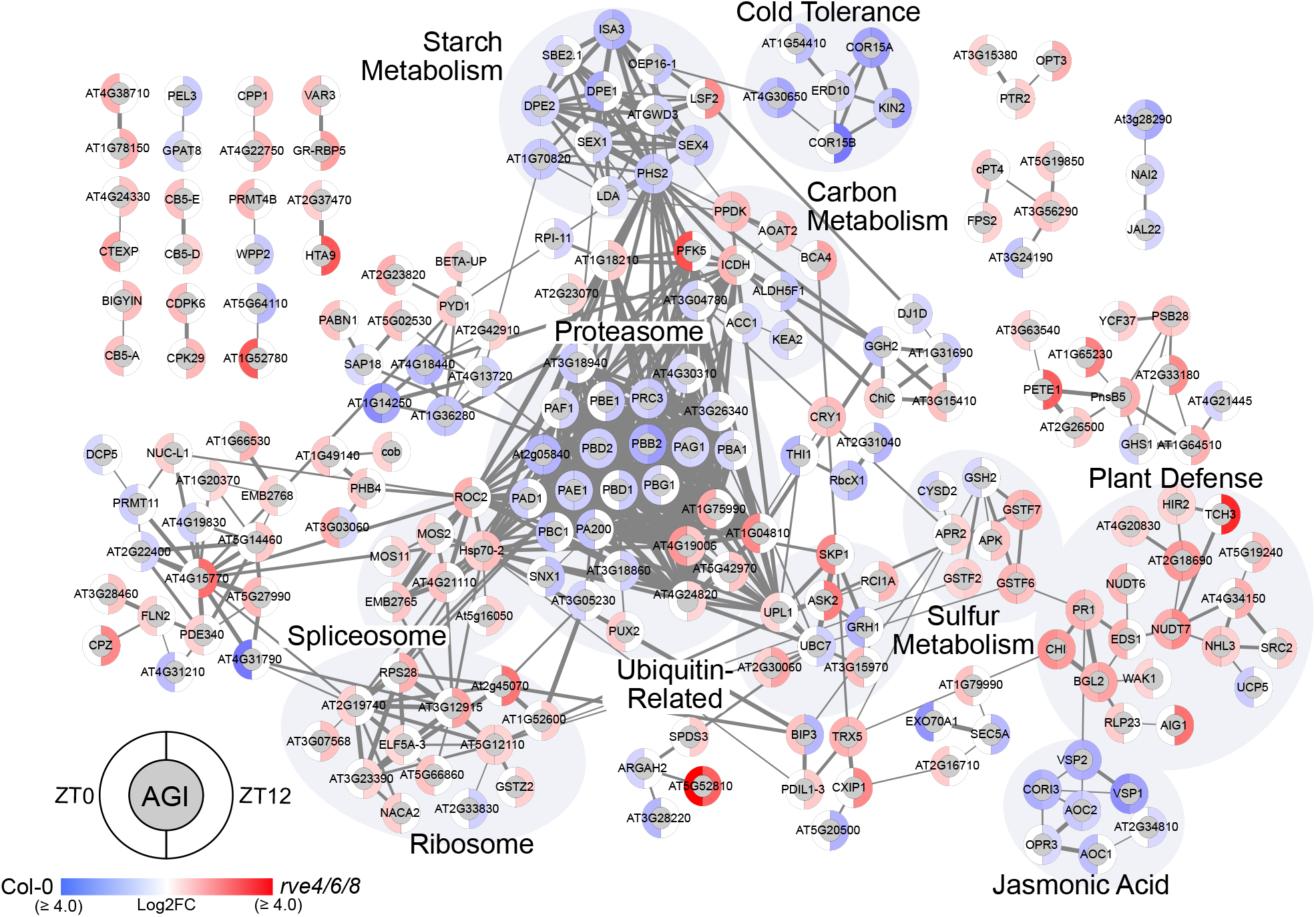
Association network analysis of significant genotypic proteome fluctuations at ZT0 and ZT12. The association network depicts significant Log_2_FC in protein abundance between *rve 4 6 8* and WT Col-0 at ZT0 and ZT12 (*q-value* ≤ 0.05; Supplemental Data 2). Networks were generated using Cytoscape and a combination of the STRING-DB and enhancedGraphics App (see Materials and Methods) using all datatypes and an edge score ≥ 0.7. Nodes with no edges ≥ 0.7 were removed. Brighter red (*rve 4 6 8*) or blue (WT Col-0) coloration indicates the relative increase in measured Log_2_FC protein abundance in that corresponding genotype. Highlighted node clusters were manually annotated using a combination of ThaleMine (https://bar.utoronto.ca/thalemine/begin.do), TAIR (https://www.arabidopsis.org/) and Cytoscape Functional Annotation.

Association network analysis also revealed that the loss of RVE8-like proteins results in major phosphoproteome perturbations, with large increases in the phosphorylation status of a number of proteins at both ZT0 and ZT12 in *rve 4 6 8* (Figure 5). Correspondingly, we observed abundance changes in protein kinases (PKs) CALCIUM DEPENDENT PROTEIN KINASE 3 (CDPK6/CPK3; AT4G23650), 29 (CPK29; AT1G76040), CYTOPLASMIC TRNA EXPORT PROTEIN (CTEXP; AT2G40730), WALL ASSOCIATED KINASE 1 (WAK1; AT1G21250), amongst others (Figure 4), in addition to phosphorylation changes on 23 protein kinase superfamily proteins including CPK1 (Figure 5). Overall, we observed phosphorylation changes on proteins involved in similar processes and protein complexes as those found in our quantitative proteome data including multiple spliceosomal- and ribosome- complex proteins, ubiquitin-related proteins and primary metabolic enzymes (Figure 5). Exclusive to our phosphoproteome data however, are perturbations in the phosphorylation status of a number of nuclear- and secretion-related proteins along with extensive changes in proteins related to light harvesting, photosynthesis and light signaling, such as CAB1 (AT1G29930), PHOT2 (AT5G58140) and HY5 (AT5G11260).

**Figure 5:**
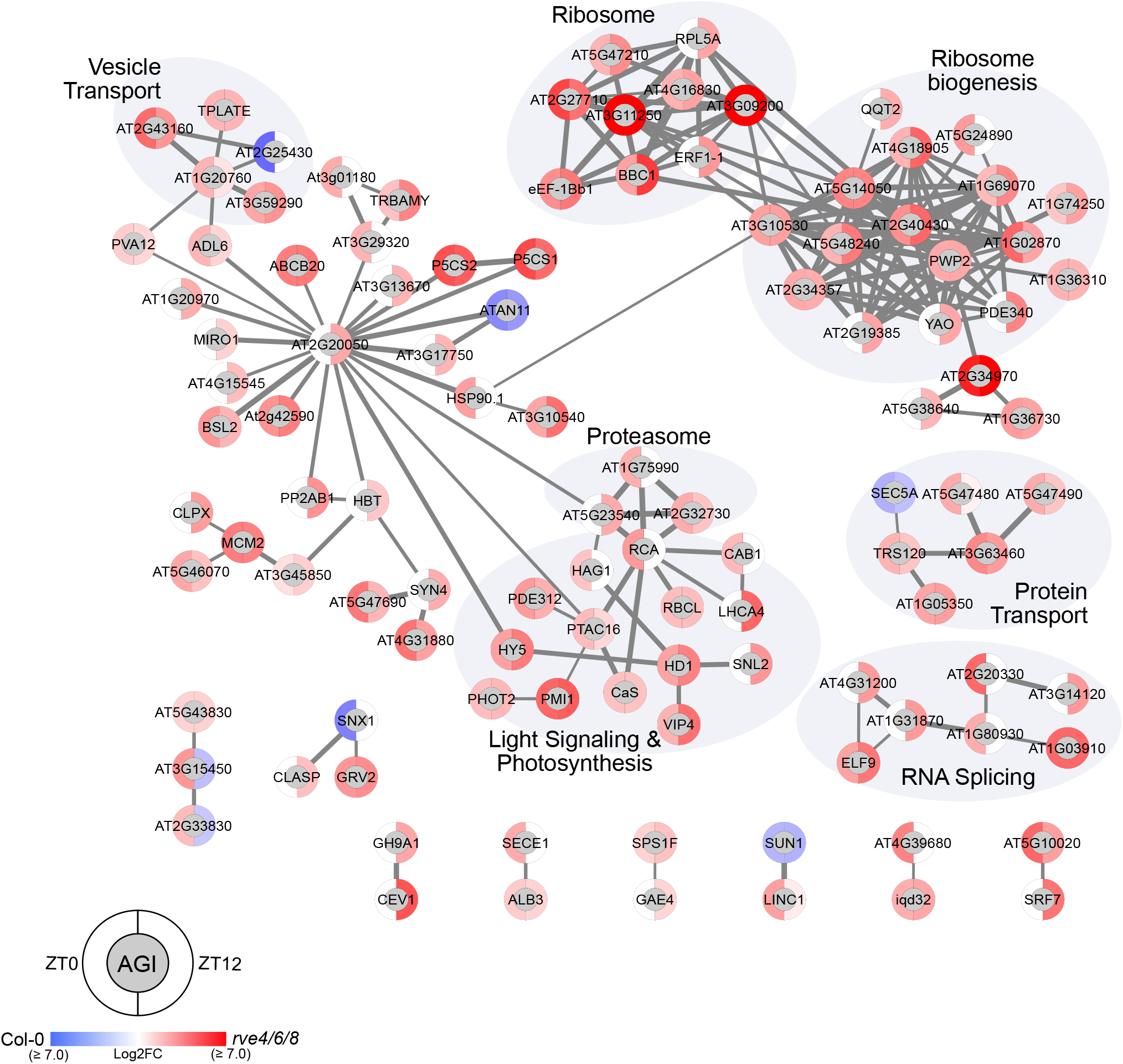
Association network analysis of significant genotypic phosphoproteome fluctuations at ZT0 and ZT12. The association network depicts significant median Log_2_FC phosphoproteome changes between *rve 4 6 8* and WT Col-0 at ZT0 and ZT12 (*q-value* ≤ 0.05; Supplemental Data 3). Networks were generated using Cytoscape and a combination of the STRING-DB and enhancedGraphics App (see materials and methods) using all datatypes and an edge score ≥ 0.7. Nodes with no edges ≥ 0.7 or only edges ≤ 0.6 were removed. Brighter red (*rve 4 6 8*) or blue (WT Col-0) coloration indicates the relative increase in measured Log_2_FC protein abundance in that corresponding genotype. Highlighted node clusters were manually annotated using a combination of ThaleMine (https://bar.utoronto.ca/thalemine/begin.do), TAIR (https://www.arabidopsis.org/) and Cytoscape Functional Annotation.

Next, we performed a phospho-motif enrichment analysis using our significantly changing phosphopeptides to identify potential PKs regulated by RVE8-like proteins. We found 5 enriched phosphorylation motifs relating to MITOGEN ACTIVATED PROTEIN KINASES (MAPKs), CASEIN KINASE II (CKII) and CALCIUM DEPENDENT PROTEIN KINASES (CDPK/CPKs) (Supplemental Data 7). Using the phosphoproteins containing these motifs, we then created a STRING-DB association network (minimum edge score ≥ 0.7) depicting the phosphorylated substrates of these PKs found in our study to resolve potential functional relationships as well as intersections between different PKs (Supplemental Figure 3). Again, a stringent minimum edge score was used to resolve robust network connections, however here, all phosphoproteins recognized as having one of these 5 motifs is shown. This analysis resolved a number of networks, including: 26S proteasomal complex proteins - MAPK, ribosome - CKII / CDPK/CPK, spliceosome - CDPK/CPK, RNA processing - MAPK and golgi-ER secretion - CDPK/CPK relationships (Supplemental Figure 3), collectively providing new cell signaling – biological process connections for further characterization.

### Wide ranging primary metabolite changes highlight altered metabolism in *rve 4 6 8* plants

As multiple proteins involved in metabolism demonstrated changes at the proteome and phosphoproteome level, we next examined diel changes in *rve 4 6 8* metabolism compared to WT Col-0 at ZT0, 6, 12 and 18 using gas chromatography mass spectrometry (Figure 6; Supplemental Data 8). Here, we found dynamic changes (*rve 4 6 8* vs. WT Col-0) in amino acids (AA), organic acids (OA), fatty acids (FA) and sugars, amongst others, over the course of the day, with the largest perturbations occurring around ZT0 - ZT6. These diel changes are highlighted by large increases in phenylalanine (AA), glutamic acid (AA), shikimic acid (OA), sinapinic acid (OA), α-linolenic acid (FA), sucrose (sugar), myo-inositol (other), 5-oxoproline (other) and phytol (other) at ZT0, along with an increase in mannose (sugar) staring at ZT6. Further, we observe that *rve 4 6 8* plants maintain a consistent increase in glyceric acid (OA) and fumaric acid (OA) coupled with a consistent decrease in serine (AA) and threonine (AA) across all four ZT time-points harvested.

**Figure 6:**
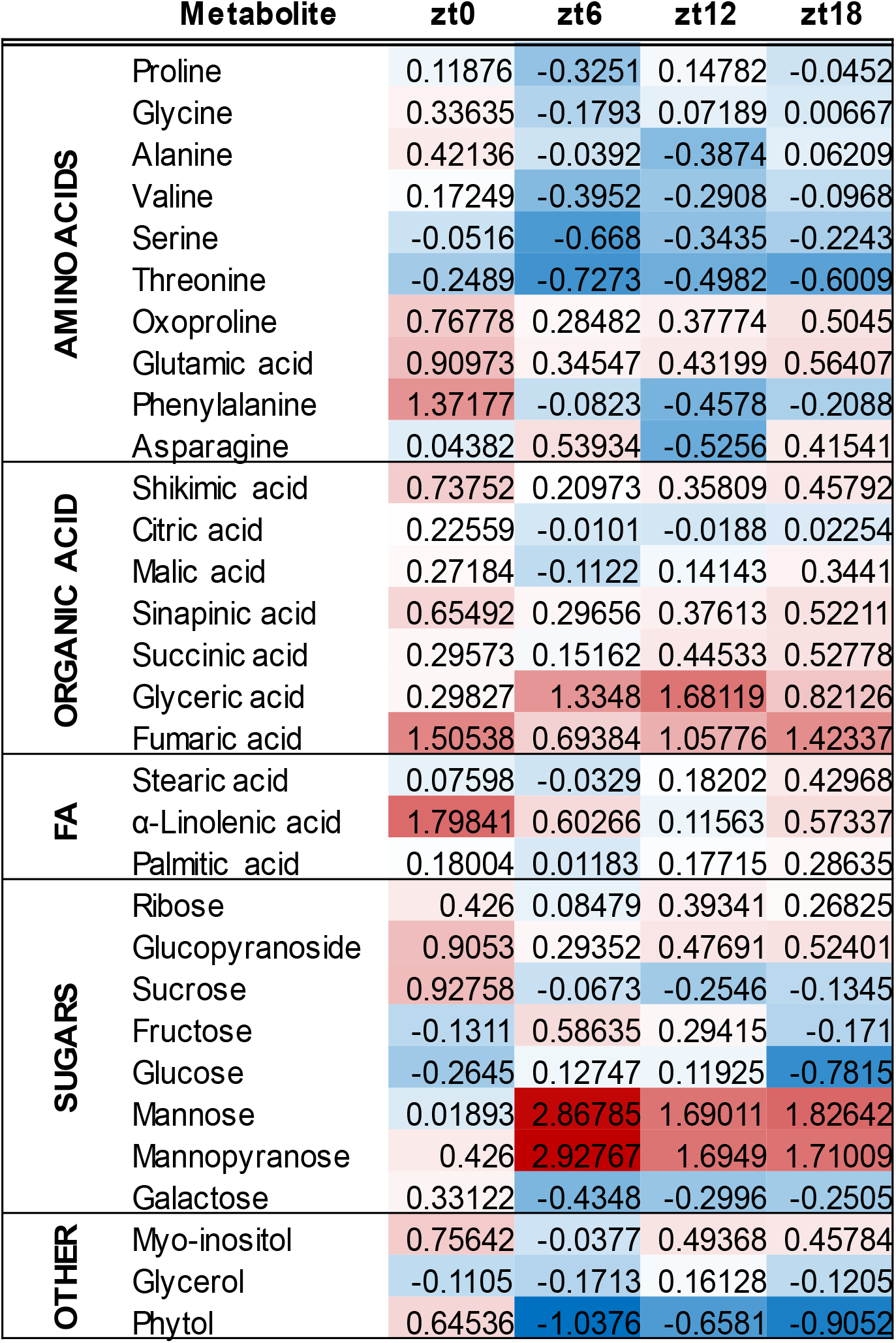
Relative Log_2_FC fold-change change in diel metabolite levels. Rosette tissue (28 d post-imbibition) was sampled at ZT0, ZT6, ZT12 and ZT18 as outlined in the Materials and Methods. Brighter red (*rve 4 6 8*) or blue (WT Col-0) coloration indicates the relative increase in measured Log_2_FC metabolite abundance in that corresponding genotype.

### *rve 4 6 8* plants possess constitutively reduced proteasome activity

The enrichment of proteolysis and proteasome related GO categories in our proteomics dataset, coupled with an extensive cluster of proteasome-related proteins in our STRING-DB association network, prompted us to examine whether the *rve 4 6 8* mutant showed differences in proteasome function compared to WT Col-0. We therefore carried out a proteasome activity assay using a 7-Amino-4-methylcoumarin (AMC) flurophore tagged LLVY peptide substrate (Üstün et al., 2017) and used the well characterized MG132 proteasome inhibitor as a negative control to validate the assay. Proteasome activity was measured in crude extracts of 8 independent wild-type and *rve 4 6 8* rosettes harvested at both ZT23 and ZT11. Our results show a significant reduction in proteasome activity in the *rve 4 6 8* mutant compared to wild-type (adjusted *p-value* ≤ 0.05; Bonferroni & Dunn multiple testing correction) (Figure 7A). Further, plants with impaired proteasome activity are also found to display a hypersensitivity to chemical proteasome inhibition (Gladman et al., 2016; Han et al., 2019). Correspondingly, when grown on plates containing 50μM of the MG132 proteasome inhibitor, *rve 4 6 8* seedlings possessed impaired growth, similar to that previously observed in mutants of both the core 20S proteasome and the NAC transcription factors that activate the proteasome stress regulon (Figure 7B, Supplemental Figure 4) (Gladman et al., 2016; Han et al., 2019).

**Figure 7:**
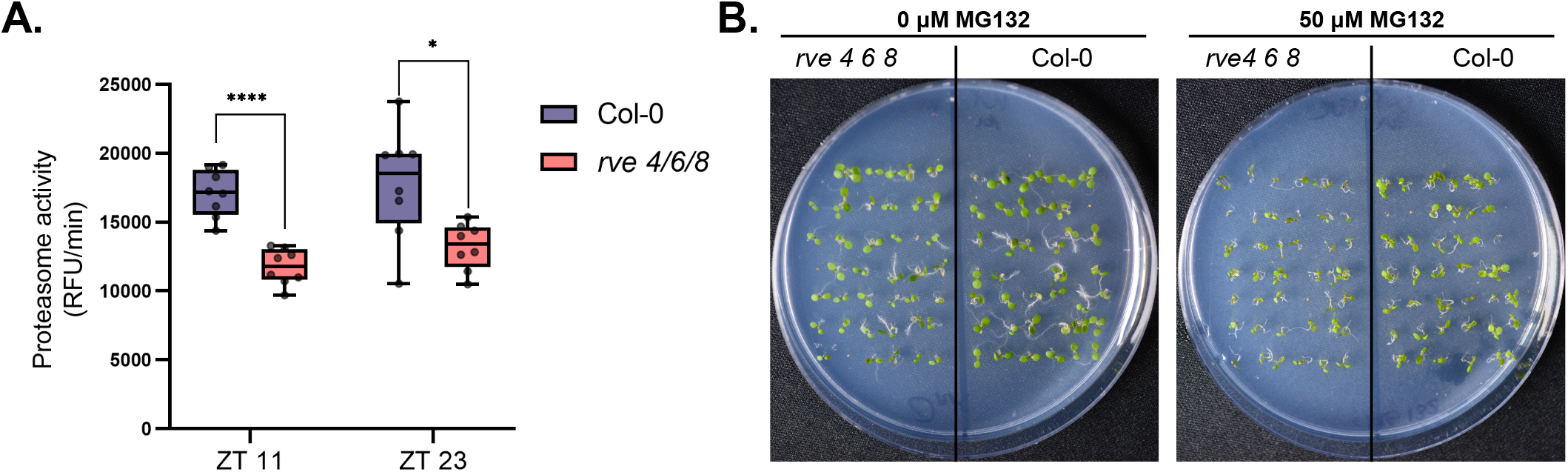
Proteasome function is reduced in *rve 4 6 8*. **(A)** Proteasome activity in relative fluorescence units per minute (RFU/min) measured in crude extracts of three week old *rve 4 6 8* and WT Col-0 plants. (**** Bonferroni adjusted *p-value* <0.0001; * Bonferroni adjusted *p-value* < 0.05) (**B**) Seedling growth experiments with proteasome inhibitor treatment (50 μM MG132). Seedlings were germinated, stratified, and then grown for 6 d under 12:12 LD conditions. A total of six replicate plates per condition were assayed (Additional plates depicted in Supplemental Figure 4).

## DISCUSSION

### Multi-omics analysis reveals altered carbohydrate metabolism in *rve 4 6 8* plants

Through our combined use of phenomics, transcriptomics, proteomics and metabolomics, we found that *rve 4 6 8* plants possess a starch excess phenotype at ZT0 that relates to altered both gene expression and protein abundance changes in starch biosynthesis and degradation enzymes. Initial examination of the transcriptional changes of select genes saw significant decreases in *GBSS1* at ZT0 and *SEX4* at ZT12 in *rve 4 6 8* relative to WT Col-0, respectively. Subsequent proteomic analysis then found extensive decreases in nearly all facets of the starch degradation machinery at either ZT0 and/or ZT12 including decreases in the abundance of SEX1 (AT1G10760), SEX4, GWD3 (AT5G26570), ISA3 (AT4G09020), PHS2 (AT3G46970), LDA (AT5G04360) and DPE2 (AT2G40840) (Streb and Zeeman, 2012). Correspondingly, our metabolite data demonstrated decreased glucose and sucrose levels at ZT6, ZT12 and ZT18, which is consistent with altered carbon metabolism related to excess starch levels at ZT0. Plants deficient in core clock proteins PRR7 and PRR9 (*prr7 prr9*), which also maintain a long circadian period, have also been shown to possess a subtle starch excess phenotype (Chew et al., 2017; Flis et al., 2019); however, unlike *prr7 prr9* plants, *rve 4 6 8* plants seem to asymmetrically possess excess starch at ZT0 only. The mutant of disproportionating enzyme (DPE1; AT5G64860), a key starch degradation protein, however, possesses a similar phenotype whereby starch levels are in excess at dawn but accumulate closer to wild-type plants by dusk (Critchley et al, 2001). We also see DPE1 asymmetrically down-regulated in protein abundance at ZT0, suggesting a connection to RVE8-like proteins. Hence, future work connecting RVE8-like proteins to DPE1 and other starch degradation enzymes is certainly of interest.

Interestingly, we see very few PTM-level changes occurring on starch-related enzymes in *rve 4 6 8* plants despite many of these enzymes being previously found to be phosphorylated (PhosPhat; http://phosphat.uni-hohenheim.de/; (Heazlewood et al., 2008)). Unlike the extensive protein abundance changes observed, only two starch-related enzymes were found to have a significant change in their phosphorylation status. These were STARCH SYNTHASE 2 (SS2; AT3G01180) and BETA-AMYLASE 1 (BAM1; AT3G23920), of which both exhibited an increase in their phosphorylation status at ZT0 and ZT12 in *rve 4 6 8* relative to WT Col-0. The role(s) of protein phosphorylation in regulating starch-related enzyme complexes has been well defined in sink tissues (Tetlow et al., 2004), but remains poorly understood in source tissues (e.g. leaves), where starch is produced and degraded on a daily basis (Smith and Zeeman, 2020). Recent biochemical characterization of SS2 phosphorylation using phospho-ablative SS2 mutants (S63A and S65A) found no enzymatic or protein complex consequences to the loss of SS2 phosphorylation despite being phosphorylated by CKII; (Patterson et al., 2018)). Our data finds SS2 to be phosphorylated at Ser65; part of a canonical CKII phosphorylation motif (*pS*[D/E][D/E][D/E]) (Purzner et al., 2018), exclusively at ZT0 in the *rve 4 6 8* mutant. We also observe BAM1 to be phosphorylated, but at a *pS*P MAPK-like motif (Purzner et al., 2018). There are no currently defined roles for BAM1 phosphorylation despite its key role in regulating plant growth, promoting stomatal opening and facilitating starch degradation under osmotic stress in the light (Thalmann et al., 2016).

Of the carbohydrate-related changes we found in *rve 4 6 8*, the consistent and substantial increase in mannose, a multifunctional glucose analog, was particularly interesting. Previous investigations of the function of mannose have found that it can: 1) stimulate stomatal closure (Hei et al., 2018), 2) be incorporated into glycolysis through fructose 6-phosphate via hexokinase and phosphomannose isomerase (Sharma et al., 2014), 3) protect against oxidative damage, 4) impact flowering time through ascorbate production (Kotchoni et al., 2009) and 5) contribute to cell wall biosynthesis and protein glycosylation (Qi et al., 2017) - all molecular changes that relate to the phenotypes displayed by *rve 4 6 8*. Interestingly, mannose has a high affinity for hexokinase (Herold and Lewis, 1977), leading to the induction of glucose-like signaling responses such as stomatal closure (Hei et al., 2018) and bud elongation in *Arabidopsis*, pea and rose (Barbier et al., 2021). It is possible that a similar mechanism contributes to the elongated rosette leaf morphology observed in *rve 4 6 8,* both in this study, and others (Gray et al., 2017).

### RVE8-like protein regulation of TOC1 may control the production of fumarate and other organic acids

*Arabidopsis* plants over-expressing RVE8 have previously been shown to stimulate the expression of *TOC1* (Hsu et al., 2013) and *toc1-2* plants have been found to possess increased levels of fumarate at ZT7 and Z19 (Cervela-Cardona et al., 2021). In this context, our finding that *rve 4 6 8* plants possess both reduced *TOC1* gene expression at ZT12 and increased fumarate levels throughout the diel cycle, suggests RVE8-like proteins impact fumarate metabolism via TOC1. Further, this finding aligns with our observation that multiple other mitochondrial organic acids are, to varying degrees, impacted by RVE8-like proteins, including malate, citrate and succinate. Change in these organic acids align with the previous findings that *prr5 prr7 prr9* mutants have increased fumarate, malate, citrate and succinate stemming from the down-regulation of TCA-cycle genes (Fukushima et al., 2009). At the protein-level however, we find no significant change in FUMARASE 1 (FUM1; AT2G47510), FUMARASE 2 (FUM2; AT5G50950), MITOCHONDRIAL MALATE DEHYDROGENASE 1 (mMDH1; AT1G53240), mMDH2 (AT3G15020), CITRATE SYNTHASE 2 (CSY2; AT3G58750), CSY3 (AT2G42790), SUCCINATE DEHYDROGENASE 1-1 (SDH1-1; AT5G66760), SDH5 (AT1G47420) and SDH6 (AT1G08480) protein abundance and/or PTM levels in *rve 4 6 8* versus WT Col-0 at either ZT0 or ZT12. This indicates that RVE8-like protein control of fumarate production is not direct, but likely through the previously identified TOC1-FUM2 mechanism (Cervela-Cardona et al., 2021). It also suggests that if diel protein-level regulation of mitochondrial TCA enzymes is occurring, it manifests through alternative means, such as a production-degradation mechanism that maintains steady state levels of TCA enzymes or through PTMs other than phosphorylation. While our current dataset is unable to directly answer these possibilities, previous work examining mitochondrial protein turnover has implicated FTSH, LON and CLPXP proteases (Li et al., 2017; Huang et al., 2020), of which we observe FTSH11 (AT5G53170) and CLPX (AT5G53350), two putatively mitochondrial targeted proteases, to possess an increase in their phosphorylation status (Heazlewood et al., 2004; Senkler et al., 2017). Ultimately how TCA enzymes are coordinately regulated at the transcript, protein and/or PTM-levels, in addition to possible feed-forward or feed-back regulation by metabolites, requires targeted analyses that are beyond the scope of this study.

### RVE8-like proteins are constitutive regulators of the proteasome

Both GO enrichment and STRING-DB association network analysis of differentially expressed proteins in *rve 4 6 8* at ZT0 and ZT12 revealed substantial protein-level changes in the abundance of a number of proteasome complex subunits, coupled with a limited corresponding change in phosphorylation status. In plants, connections between the circadian clock and E3 ubiquitin ligases targeting proteins for proteasomal degradation through K63 polyubiquitylation have emerged, indicating that diel protein turnover is a circadian controlled phenomenon (Feke et al., 2019; Feke et al., 2020; Zhu et al., 2020a). Further, using N^15^ labeling methods, efforts made to quantify protein turnover rates have demonstrated that diel protein turnover involves a multitude of proteins that are processed at variable rates over a wide-ranging time frame (Li et al., 2017; Huang et al., 2020). To date however, no connections between the circadian clock and the direct regulation of the proteasome complex have been uncovered.

The mature 26S proteasome is comprised of a 20S core protease (CP) which maintains non-specific ubiquitin-independent proteolytic activity and 19S regulatory particle (RP) subunits that determine specificity to ubiquitylated proteins (Sadanandom et al., 2012). We observe a reduction in 12 out of the 14 CP subunits, including 5 alpha and 7 beta ring subunits, in the *rve 4 6 8* mutants. In contrast, we see an increase in the abundance of 4 RP subunits, as well as CSN4, a member of the COP9 signalosome, a conserved regulator of ubiquitin-dependent proteolysis and a key player in photomorphogenesis (Qin et al., 2020). Studies investigating proteasome subunits in Arabidopsis have shown that reduced CP proteolytic function is associated with cell expansion (Kurepa et al., 2009a) and that one member of the 19S RP subunit, RPN12a, is involved in cytokinin-mediated signaling for cell division (Smalle et al., 2002). This aligns with our GO enrichment of cell cycle and cell division biological process amongst proteins exhibiting a significant change in abundance as well as our observed increase in cytokinin response factor ARR7. Further, these findings are consistent with a hypothesis that proper balancing of the 20S and 19S subunits of the proteasome is key for maintaining growth and cell division (Kurepa et al., 2009b), and that alterations in the levels of these subunits in *rve 4 6 8*, at least partially, explain its growth and large cell size phenotypes.

In mice, connections between the circadian clock and proteasomal degradation of CRYPTOCHROME 1 (CRY1) have also been defined, with enhanced CRY1 degradation associated with elevated glucose levels and a loss of core clock regulation (Toledo et al., 2018). In plants, CRY proteins entrain the clock (Devlin and Kay, 2000), with *cry1* mutants having longer circadian periods under blue light (Somers et al., 1998). Interestingly, in *rve 4 6 8* plants, which similarly possess a long circadian period that increases with blue light fluence (Gray et al., 2017), we observe increased CRY1 protein levels at both ZT0 (Log_2_FC = 1.05) and ZT12 (Log_2_FC = 0.81), coupled with a starch excess phenotype and wide-ranging carbohydrate metabolism perturbations, indicating a potentially similar circadian clock intersection between CRY1 and primary carbohydrate metabolism that appears to involve RVE8-like proteins. Additionally, the diel amino acid perturbations measured in *rve 4 6 8* plants may also correspond to RVE8-like proteins impacting the diel plant cell environment through the regulation of proteasome subunits and protein turnover.

Ultimately, a connection between the circadian clock and diel protein turnover is logical given the consistent lack of coordinately fluctuating diel transcripts and proteins in *Arabidopsis* across multiple experiments (Baerenfaller et al., 2012; Graf et al., 2017; Mehta et al., 2021; Uhrig et al., 2021). Overall, our results implicate RVE8-like proteins as being involved in diel protein turnover, adding important context to the well-known disconnect between circadian transcript and protein abundancies. Future assessment of core circadian clock mutants for changes in protein turnover rates may help resolve the degree to which diel protein turnover contributes to daily protein-level plant cell regulation.

### Proteome changes in *rve 4 6 8* plants suggest RVE8-like proteins may modulate adaptive plant stress responses

The enrichment of GO categories pertaining to Abscisic acid in the *rve 4 6 8* proteome, provides protein-level evidence that RVE8-like proteins may modulate adaptation to drought-like stresses (e.g. osmotic, salinity, and cold). A potential role for RVE8-like proteins in drought-like stress response(s) is supported by the measured increase in mannose-levels, which is known to influence stomatal conductance. Further, α-linolenic acid, and its derivatives, are known to increase upon drought and salinity stress to mitigate associated cell membrane issues by remodeling cell membranes to increase their fluidity (Zhang et al., 2005; Upchurch, 2008; Torres-Franklin et al., 2009; He and Ding, 2020). Here, this seems to center on FATTY ACID DESATURASE 7 (FAD7; AT3G11170), which utilizes linoleic acid to produce α-linolenic acid (Gibson et al., 1994), and is significantly more abundant in *rve 4 6 8* plants at ZT0, corresponding to peak levels of α-linolenic acid.

In addition to augmenting membrane fluidity, α-linolenic acid can also fuel the production of jasmonic acid (JA) (He and Ding, 2020). Interestingly, we find that critical jasmonic acid biosynthetic enzymes AOC1, AOC2 and 12-OXOPHYTODIENOATE REDUCTASE 3 (OPR3; AT2G06050) are significantly more abundant in WT Col-0 relative to *rve 4 6 8*, suggesting that RVE8-like protein deficient plants may possess susceptibility to certain biotic stresses. A diel connection between RVE8-like proteins and JA biosynthetic enzymes is further supported by the altered diel phosphorylation of the peroxisomal ABC transporter COMATOSE (CTS; AT4G39850), which imports JA precursor *cis*-12-oxophytodienoic acid that is then utilized by OPR3 (Wasternack and Hause, 2019), which peaks in protein abundance at ZT12.

Further support for a role of RVE8-like proteins in drought-like stress response(s) is the increased abundance of key sulfur assimilating enzymes APR2 (AT1G62180) and APK (AT2G14750) in *rve 4 6 8* plants. These proteins are critical for the production of ABA and glutathione (GSH), in addition to a number of other critical plant metabolites (Aarabi et al., 2020). Correspondingly, we see an increase in a number of glutathione transferases (GSTs); GSTF2, GSTF6 (AT1G02930) and GSTF7 (AT1G02920), which are a subclass of the plant-specific Phi-family of GSTs implicated in mitigating oxidative (GSTF2) and salt (GSTF6/7) stress, respectively (Lee et al., 2014; Seok et al., 2020). In previous proteomic and phosphoproteomic analyses of persistent osmotic and salt-stress, increased abundance of sulfur metabolic enzymes (Rodriguez et al., 2021), indicates a potential connection between sulfur metabolism / assimilation and drought-stress tolerance. In maize, the application of sulfur-supplemented fertilizers to drought stressed plants was shown to mitigate the effects of drought (Usmani et al., 2020).

RVE8-like proteins were most recently implicated in the direct transcriptional activation of DEHYDRATION RESPONSE ELEMENT B (DREB) expression under cold-stress (Kidokoro et al., 2021). DREB transcription factors serve as master controllers of cold-induced gene expression, regulating COLD REGULATED (COR) genes to enhance freezing tolerance (Jaglo-Ottosen et al., 1998). Correspondingly, RVE8-like protein deficient plants (*rve 4 8*) possessed diminished cold tolerance (Kidokoro et al., 2021). Interestingly, our proteomics data resolved a cluster of cold-stress related proteins that are reduced in abundance at ZT12 / ED under normal growth temperatures, suggesting that some cold response proteins are diel regulated, likely possessing additional diel roles beyond cold-stress response (Maszkowska et al., 2018; Zhu et al., 2020b). These include: chloroplast targeted COR15A (AT2G42540) and COR15B (AT2G42530) (Artus et al., 1996) as well as cytoplasmic-targeted KIN2/COR6.6 (AT5G15970), ERD10 (AT1G20450), AT1G54410 and AT4G30650. In conjunction with our analyses, these findings directly connect RVE8-like proteins to observable proteome-level changes in genes associated with drought-related stresses.

### Protein kinases impacted by RVE8-like proteins

In addition to catalyzing the phosphorylation of target proteins, protein kinases (PKs) themselves are often phosphorylated, resulting in changes in their activity, function or localization, forming a complex signaling web that can have far-reaching ramifications for plant cell function. Our combined quantitative proteomic and phosphoproteomic analysis uncovered a number of PKs that change in either diel abundance or phosphorylation status in response to the loss of RVE8-like proteins. Of particular interest is the increased abundance of CASEIN KINASE II (CKIIα; AT2G23070), CALCIUM DEPENDENT PROTEIN KINASE 6 (CDPK6/CPK3; AT4G23650) and WALL ASSOCIATED KINASE 1 (WAK1; AT1G21250) and the phosphorylation of CPK1 (AT5G04870), as these PKs have been implicated in multiple plant cell processes related to *rve 4 6 8* phenotypes as well as osmotic- and salt-stress tolerance.

CKII is a well-known clock associated kinase that directly affects the stability of core clock component CCA1 in *Arabidopsis* (Sugano et al., 1998; Lu et al., 2011). The role of CKII in modulating the circadian clock is conserved across eukaryotes, with both mammalian (Tamaru et al., 2009) and Drosophila (Akten et al., 2003) circadian clocks being directly impacted by CKII. Most recently, multiple CASEIN KINASE 1 family kinases were discovered to phosphorylate PRR5 and TOC1 leading to their degradation (Uehara et al., 2019). Our finding that CKII levels are elevated in *rve 4 6 8* at ZT12 hints at the possibility of RVE8-like proteins similarly directing CKII-mediated phospho-regulation of circadian function. Beyond the clock, CKII has been implicated in phosphorylating numerous light signaling proteins, such as HY5 (Hardtke et al., 2002), which we see here possessing an increase in its phosphorylation status in *rve 4 6 8* plants at Ser36 at both ZT0 and ZT12. Recently, HY5 phosphorylation at Ser36 was found to be phosphorylated by SUPPRESSOR OF PHYTOCHROME A-105 (SPA) kinases (Wang et al., 2021), suggesting that perhaps other CKII-motif containing proteins resolved here may also represent SPA targets. Lastly, CKII is also characterized as phosphorylating ABA / abiotic stress induced pathway proteins (Mulekar and Huq, 2014), which aligns with our enrichment of ABA-related GO terms in our phosphoproteomic data.

CDPK6/CPK3 is a positive regulator of ABA-induced regulation of stomatal closure (Mori et al., 2006) and is induced under high salt conditions (Mehlmer et al., 2010), suggesting that its increased abundance in *rve 4 6 8* plants may convey signaling events related to osmotic- / salt-stress response. Although CDPK6/CPK3 has not been shown to directly impact the circadian clock, its connection to stomatal regulation, coupled with known diel fluctuations in calcium levels (Marti Ruiz et al., 2020), suggests a connection to the regulation of diel plant cell processes. Similar to CDPK6/CPK3 salt-stress induction, over-expression of native CPK1 resulted in increased salinity tolerance in cell culture (Veremeichik et al., 2021), reinforcing a potential link between diel CPK-mediated phosphorylation events and the ability to mitigate abiotic stress. Alternatively, WAK1 has been implicated in maintaining cell wall integrity (Leschevin et al., 2021), which has diel plant cell growth implications that may contribute to the large mesophyll cell phenotype of *rve 4 6 8* plants (Gray et al., 2017).

These results, when combined with our phospho-motif analysis that shows an enrichment in CKII and CDPK/CPK target motifs, suggest that CKIIα and CDPK6/CPK3 may be connected to a substantial portion of the observed RVE8-like dependent phosphoproteome. STRING-DB association network analysis of proteins with these enriched phospho-motifs demonstrates connections to multiple nuclear protein networks including one involving RETINOBLASTOMA-RELATED 1 (RBR; AT3G12280), a critical growth and development protein, in addition to protein translation and secretion. Collectively, these data provide novel insights into RVE8-like protein dependent phospho-networks and the corresponding kinases that may be involved.

## CONCLUSION

Our robust, multi-omic analysis of *rve 4 6 8* plants further establishes RVE8-like proteins as targets of agronomic interest by defining the proteomic and metabolic underpinnings of their observable phenotypes. Through these analyses, we found a number of connections to clock related cell processes that refine our understanding of *rve 4 6 8* growth phenotypes. This includes altered metabolism that either directly (e.g. carbohydrates and organic acids) or indirectly (e.g. precursors for growth phytohormones) drives growth outcomes. In particular, carbohydrate metabolism, for which we found a significant decrease in multiple starch degradation enzymes leading to starch excess at dawn. As well, our findings reveal that RVE8-like proteins control the 26S proteasome through altered 20S and 19S subunit abundance and phosphorylation status, which our metabolite data suggests may be tied to changes in resource pools important for growth and development (e.g. amino acids). This novel connection between the proteasome and the circadian clock through RVE8-like proteins was further supported by a reduction of proteasome activity in *rve 4 6 8* plants, something that in proteasome mutants, leads to a large cell size phenotype similar to that observed in *rve 4 6 8* plants. This suggests that RVE8-like proteins regulate the proteasome as a means of controlling cell division and growth. Lastly, our multi-omic analyses hint at potential new connections between RVE8-like proteins and abiotic stress response (e.g. osmotic, salt and/or cold – already established), which we see manifesting throughout our proteome, phosphoproteome and metabolome data. Overall, our combined use of quantitative transcriptomics, proteomics, PTMomics and metabolomics have elucidated a number of new connections between RVE8-like proteins and critical plant cell processes, laying a substantial foundation from which to build a comprehensive understanding of the roles RVE8-like proteins play in directing diel plant cell regulation.

## MATERIALS AND METHODS

### Plant Growth and Phenotyping

#### Plant Growth

Wild-type *Arabidopsis thaliana* Col-0 and *rve 4 6 8* seeds were obtained from Dr. Stacy Harmer (University of California - Davis). Seeds were sterilized in 70% (v/v) ethanol for 2 min followed by a 30% (v/v) bleach (Chlorox 7.5%) treatment for 7 min and three washes with distilled water. The seeds were then grown on 0.5x MS media and 7 g/L of agar at pH 5.8 (with KOH). All seeds were cold treated for 3 days at 4°C in the dark and exposed to light for a week before being transferred to soil (Sungro, Sunshine Mix® #1). After 21 days of light exposure, entire plants were collected for analysis. Growth chambers were equipped with a programmable Perihelion LED fixture (G2V Optics Inc.) and lined with Reflectix® to ensure a good light diffusion. Plants were grown under a 12h light and 12h dark photoperiod with a temperature of 21°C during the day and 19°C at night. Six LED types (Cree LED XPE, XPE2 XPG families) were used in the fixtures, with characteristic wavelengths of 444 nm, 630 nm, 663 nm, 737 nm, 3000 K white, and 6500 K white.

#### Phenomics measurements

Each chamber was equipped with a Raspberry Pi 3 B+ and an ArduCam Noir Camera (OV5647 1080p) with a motorized IR-CUT filter and two infrared LEDs. Pictures were taken every 15 min over 7 d (between 14 d and 21 d post seed imbibition) and were used to extract plant area and perimeter measurement using PlantCV as previously described (Gehan et al., 2017).

#### Proteasome inhibitor sensitivity seedling growth experiment

Col-0 and *rve 4 6 8* seedlings were germinated on 0.5x MS control plates and 0.5x MS plates supplemented with 50 μM MG132 (Millipore Sigma; 474787) and grown under 12:12 photoperiod conditions as described above for 6 days prior to imaging.

### Starch Iodine Staining and Enzymatic Assay

#### Starch Iodine Staining and quantification

Starch was stained and visualized in *Arabidopsis* rosettes by Lugol’s iodine stain. Entire rosettes were decolorized by incubation in 80% ethanol at 80°C for 30 minutes. Rosettes were rinse 3 times in distilled water and stained 20 min in a 5% (v/v) Lugol’s iodine solution (Fisher Scientific). After 3 rinses in distilled water, rosettes were washed for an 1h in distilled water to remove the excess of iodine. Rosette leaves were flattened between plastic sheet and excess water removed. Each rosette was analyzed with ImageJ (Abramoff et al 2004; https://imagej.nih.gov/ij/index.html) to record mean values in starch staining pixel intensity as described by Betti et al., (2016).

#### Starch Extraction and Quantification

Starch was extracted and assayed as previously described (Smith and Zeeman, 2006). Frozen rosette leaves were ground to a fine powder and homogenized in perchloric acid 8% (Ricca Chemical). The insoluble fraction was pelleted and washed 3 times in ethanol 80% and resuspended in water. Prior quantification, the samples were boiled 15 min at 95°C to gelatinize the starch and hydrolyzed with α-amylase and amyloglucosidase (Roche) to release the glucose blocks. Glucose content was then measured by spectrophotometry after a 2 enzymes reaction with hexokinase and glucose-6-phosphate dehydrogenase (Roche).

### RNA isolation and NanoString

Total RNA was isolated and purified using a modified single step TRIzol protocol (Chomczynski and Sacchi, 1987). Plants were flash frozen and homogenized with two glass beads using a Geno/Grinder® for 30s at 1500 x g. Immediately after, 1 mL of TRIzol was added to the sample, and the whole was vortexed and incubated for 10 min at room temperature. The samples were then centrifuged at 13000 x g for 10min to remove extracellular material and glass beads. The supernatant was transferred to a new tube. Subsequently, 200 μL of chloroform was added to the mixture and shaken vigorously by hand for 15 s and the samples were kept for 3 min at room temperature. The samples were centrifuged at 13000 x g for 15 min at 4°C and the supernatant thus obtained was transferred to a new tube by gentle pipetting, avoiding the interphase slurry. RNA was precipitated by adding 500 μL of 100% isopropanol and the tubes were incubated 10 min at RT and centrifuged at 13000 x g for 10 min at 4°C. After decanting isopropanol, the pellet was washed with 1 mL of 75% ethanol in nuclease-free water (IDT^TM^) and briefly centrifuged at 7500 x g for 5 min at 4°C. Total RNA was suspended in nuclease-free water and stored at 80°C for further use. RNA quantification was performed using a NanoDrop ND 1000 spectrophotometer. The RNA was treated with DNase I, Amp grade (Invitrogen) following the instructions provided by the manufacturer. The purified RNA thus obtained was reverse to cDNA using the RevertAid RT Kit (Thermo Scientific), according to manufacturer’s instructions. A minimum of 150ng of purified RNA in 7.5ul was sent to NanoString (NanoString Technologies, Seattle, USA, https://www.nanostring.com) for analysis. The NanoString probes were designed and synthesized by NanoString (Supplementary Data 1). UBC21 (AT5G25760) was used as reference gene.

### Liquid Chromatography Mass Spectrometry (LC-MS)

#### Protein Extraction

Quick-frozen rosette tissue was ground to a fine powder under liquid N_2_ using a mortar and pestle. Ground samples were aliquoted into 400 mg fractions. Aliquoted samples were then extracted at a 1:2 (w/v) ratio with a solution of 50 mM HEPES-KOH pH 8.0, 50 mM NaCl, and 4% (w/v) SDS. Samples were then vortexed and placed in a 95°C table-top shaking incubator (Eppendorf) at 1100 RPM for 15 mins, followed by an additional 15 mins shaking at room temperature. All samples were then spun at 20,000 x g for 5 min to clarify extractions, with the supernatant retained in fresh 1.5 mL Eppendorf tubes. Sample protein concentrations were measured by bicinchoninic acid (BCA) assay (23225; Thermo Scientific). Samples were then reduced with 10 mM dithiothreitol (DTT) at 95°C for 5 mins, cooled, then alkylated with 30 mM iodoacetamide (IA) for 30 min in the dark without shaking at room temperature. Subsequently, 10 mM DTT was added to each sample, followed by a quick vortex, and incubation for 10 min at room temperature without shaking. Total proteome peptide pools were then generated using a KingFisher Duo (Thermo Scientific) automated sample preparation device as outlined by Leutert et al. (2019) without deviation. Sample digestion was performed using sequencing grade trypsin (V5113; Promega), with generated peptide pools quantified by Nanodrop, normalized to 400 μg for phosphopeptide enrichment and acidified with formic acid (FA) to a final concentration of 5% (v/v) prior to being desalted using 1cc tC18 Sep Pak cartridges (WAT036820; Waters) as previously described (Uhrig et al., 2019).

#### Phosphopeptide Enrichment

SepPak eluted peptides were topped up to 65% (v/v) acetonitrile (ACN) and 5% (v/v) trifluoroacetic acid (TFA) prior to enrichment using a KingFisher Duo (Thermo Scientific) automated sample preparation device as outlined by Leutert et al. (2019). We used 500 μg of MagReSyn Ti-IMAC HP matrix (Resyn BioSciences) to enrich phosphorylated peptides. Unbound peptides (total proteome) and phosphopeptides were dried then dissolved in 3% (v/v) ACN / 0.1% (v/v) TFA, desalted using ZipTip C18 pipette tips (ZTC18S960; Millipore) as previously described (Uhrig et al., 2019). All peptides were then dried and re-suspended in 3% (v/v) ACN / 0.1% (v/v) FA immediately prior to MS analysis.

#### BoxCarDIA Mass Spectrometry

Changes in the protein abundancies was assessed using trypsin digested samples analysed using a Fusion Lumos Tribrid Orbitrap mass spectrometer (Thermo Scientific) in a data independent acquisition (DIA) mode using the BoxCarDIA method (Mehta et al., 2022). Dissolved peptides (1 μg) were injected using an Easy-nLC 1200 system (LC140; Thermo Scientific) and separated on a 50 cm Easy-Spray PepMap C18 Column (ES803A; Thermo Scientific). A spray voltage of 2.2 kV, funnel RF level of 40 and heated capillary at 300°C was deployed, with all data acquired in profile mode using positive polarity, with peptide match turned off and isotope exclusion selected. All gradients were run at 300 nL/min with analytical column temperature set to 50°C. Peptides were eluted using a segmented solvent B gradient of 0.1% (v/v) FA in 80% (v/v) ACN from 4% - 41% B (0 - 107 min). BoxCar DIA acquisition was performed as previously described (Mehta et al., 2022). MS^1^ analysis was performed by using two multiplexed targeted SIM scans of 10 BoxCar windows each, with detection performed at a resolution of 120,000 at 200 m/z and normalized AGC targets of 100% per BoxCar isolation window. Windows were custom designed as previously described (Mehta et al., 2022). An AGC target value for MS^2^ fragment spectra was set to 2000%. Twenty-eight 38.5 m/z windows were used with an overlap of 1 m/z. Resolution was set to 30,000 using a dynamic maximum injection time and a minimum number of desired points across each peak set to 6.

#### BoxCarDIA Data Analysis

All acquired BoxCar DIA data was analyzed in a library-free DIA approach using Spectronaut v14 (Biognosys AG) using default settings. Key search parameters employed include: a protein, peptide and PSM FDR of 1%, trypsin digestion with 1 missed cleavage, fixed modification including carbamidomethylation of cysteine residues and variable modifications including methionine oxidation. Data was Log_2_ transformed, globally normalized by median subtraction with significantly changing differentially abundant proteins determined and corrected for multiple comparisons (Bonferroni-corrected *p-value* ≤ 0.05; *q-value ≤ 0.05*).

#### DDA Mass Spectrometry

Changes in the protein phosphorylation status was assessed using trypsin digested samples analyzed using a Fusion Lumos Tribrid Orbitrap mass spectrometer in a data dependent acquisition (DDA) mode using the Universal method (Thermo Scientific). Dissolved phosphopeptides were analyzed as described above, using a linear gradient of solvent B (0.1% (v/v) FA in 80% (v/v) ACN): 5% to 22% B, 0 – 110 min; 22% - 35% B, 110 – 120 min; 35 - 95% B, 120 – 125 min at a flow rate of 0.3 μl/min at 50°C. Full scan MS1 spectra (350 - 2000 m/z) were acquired with a resolution of 120,000 at 200m/z with a normalized AGC Target of 125% and a maximum injection time of 50 ms. DDA MS2 were acquired in the linear ion trap using quadrupole isolation in a window of 2.5 m/z. Selected ions were HCD fragmented with 35% fragmentation energy, with the ion trap run in rapid scan mode with an AGC target of 200% and a maximum injection time of 100 ms. Precursor ions with a charge state of +2 - +7 and a signal intensity of at least 5.0e3 were selected for fragmentation. All precursor signals selected for MS/MS were dynamically excluded for 30s.

#### DDA Data Analysis

All acquired DDA phosphoproteomic data was analyzed using MaxQuant 1.6.14.0 (http://www.maxquant.org/; (Cox and Mann, 2008)) with the following parameters. A protein, peptide and PSM FDR of 1%, trypsin digestion with 1 missed cleavage, fixed modification including carbamidomethylation of cysteine residues and variable modifications including methionine oxidation and phosphorylated serine, threonine and tyrosine. Data was then imported into Perseus version 1.6.14.0 (Tyanova et al., 2016) for further analysis. This involved the removal of reverse hits and contaminants, log_2_-transformation, assembly into treatment groups and filtering based on the presence of measured data in at least 2/3 replicates of at least one group. Data were then median normalized, followed by imputation of missing values based on the normal distribution function set to default parameters. All PTM analyses utilized a PTM site localization score threshold ≥ 0.75. Significantly changing differentially abundant phosphopeptides were determined by ANOVA and corrected for multiple comparisons (Benjamini-Hochberg corrected *p-value* ≤ 0.05; *q-value*).

All proteomic and phosphoproteomic raw data and search parameters have been uploaded to ProteomeXchange (http://www.proteomexchange.org/) via the Proteomics IDEntification Database (PRIDE; https://www.ebi.ac.uk/pride/). Project Accession: PXD029234.

### Gas Chromatography Mass Spectrometry (GC-MS)

#### Metabolite Extraction and preparation

Metabolite extraction and preparation was performed with modifications as previously described (Hill and Roessner, 2013; Liu et al., 2016). Tissue was harvested and directly flash frozen in liquid nitrogen. Samples of 100 mg (+/− 1 mg) of pulverized tissue were prepared and homogenized in 700 μl of iced-cold methanol (80% v/v). Internal standard 25 μl ribitol at 0.4 mg.ml^−1^ in water. Samples were incubated 2 h at 4°C with shaking and then 15 min at 70°C at 850 rpm in a Thermomixer. Tubes were centrifuged 30 min at 12000 rpm and the supernatants were transferred in new tubes. Polar and non-polar phases were separated by the addition of 700 μl of water and 350 μl of chloroform, then vortexed thoroughly and centrifuged for 15 min at 5000 rpm. The upper methanol/ water phase (150 μl) was transferred to a new tube and dried in a vacuum centrifuge at RT. Samples were derivatized with 100 μl of methoxylamine hydrochloride (20 mg.ml^−1^ in pyridine) for 90 min at 30°C at 850 rpm in thermomixer and followed by incubation with 100 μL of N,O-bis(trimethylsilyl)trifluoroacetamide (BSTFA) at 80°C during 30 min with shaking at 850 rpm in thermomixer.

#### GC-MS analysis

Finally, samples were injected in splitless mode and analyzed using a 7890A gas chromatograph coupled to a 5975C quadrupole mass detector (Agilent Technologies, Palo Alto, CA, USA). In the same manner 1 μl of retention time standard mixture Supelco C7–C40 saturated alkanes standard (1,000 μg.ml^−1^ of each component in hexane) diluted 100 fold (10 μg.ml^−1^ final concentration) was injected and analyzed. Alkanes were dissolved in pyridine at 0.22 mg.ml^−1^ final concentration. Chromatic separation was done with a DB-5MS capillary column (30 m × 0.25 mm × 0.25 μm; Agilent J&W Scientific, Folsom, CA, USA). Inlet temperature was set at 280°C. Initial GC Oven temperature was set to 80°C and held for 2 min after injection then GC oven temperature was raised to 300°C at 7°C min^−1^, and finally held at 300°C for 10 min. Injection and ion source temperatures were adjusted to 300°C and 200°C, respectively with a solvent delay of 5 min. The carrier gas (Helium) flow rate was set to 1 ml.min^−1^. The detector was operated in EI mode at 70 eV and in full scan mode (m/z 33–600).

#### Metabolite identification and quantification

Compounds were identified by mass spectral and retention time index matching to the mass spectra of the National Institute of Standards and Technology library (NIST20, https://www.nist.gov/) and the Golm Metabolome Database (GMD, http://gmd.mpimp-golm.mpg.de/). Metabolites quantification was performed using MassHunter Software from Agilent. Peaks were integrated and after blank subtraction were normalized by dividing them by the peak area of the internal standard ribitol and by the sample weight.

### Bioinformatics Analysis and Data Visualization

Visualizations were made using *R* version 3.6.1 and R package *ggplots* in combination with Affinity Designer (https://affinity.serif.com/en-us/designer/; ver. 1.9.1.179). All data was plotted using GraphPad - Prism (ver. 8; https://www.graphpad.com/scientific-software/prism/). Networks were created in Cytoscape (ver 3.7.1; https://cytoscape.org/), using the STRING-DB (https://string-db.org/) and enhancedGrahphics (ver. 1.5.4; https://apps.cytoscape.org/apps/enhancedgraphics) cytoscape-apps. All association networks were built without data sub-selection and using an edge threshold ≥ 0.7). Gene Ontology Enrichments were performed using TheOntologizer (http://ontologizer.de/) using all identified proteins as the background for enrichment analyses (*q-value* ≤ 0.05), while phosphorylation motif enrichment analysis was performed using Motif-X (Chou and Schwartz, 2011) on the MoMo MEME suite (https://meme-suite.org/meme/) as previously described (Mehta et al., 2021). Subcellular localization information for proteins was obtained from SUBA4 (https://suba.live/).

### Proteasome activity assay

Proteasome activity was measured using the fluorogenic substrate N-Succinyl-Leu-Leu-Val-Tyr-7-Amido-4-Methylcoumarin (Millipore Sigma; S6510) as described Üstün et al. (2017). Briefly, 100 μg of rosette tissue of 3 week old plants was extracted with 250 μL of extraction buffer (2 mM ATP, 50 mM HEPES-KOH (adjusted to pH 7.2 with 1 N KOH); 2 mM DTT; 0.25 M sucrose). After clarifying by centrifugation at 18,000 g for 15 minutes, total protein content in the extracts was quantified with a Bradford assay (Bio-Rad; 5000006). The assay was performed using 50 μg of total protein extract and 300 μM of substrate in assay buffer (100mM HEPES-KOH pH 7.8, 5mM MgCl_2_, 10 mM KCl, and 2 mM ATP) at 30°C. Fluorescence was measured at 1.5 minute intervals with an excitation at 360nm and emission wavelength at 460nm for a period of 2 hours after an initial 2 hour incubation. Activity was reported as Relative Fluorescence Units/min based on the slope of the linear portion of the emission curve (MeanV in the Gen5 software).

## ACKNOWLEDGEMENTS

The authors would like to thank Dr. Stacey Harmer for providing *rve 4 6 8* and corresponding WT Col-0 seeds as well as Dr. Pascal Schläpfer for assistance with the MoMo MEME suite. We would also like to thank G2V Optics Inc. for providing their LED lighting system. As well, we would like to thank Jack Moore of the Alberta Mass Spectrometry and Proteomics Facility as well as Broderick Wood, Jacob Blazusiak and Mohamad Jamaleddine of the University of Alberta Faculty of Science Research IT team for their technical assistance.

## SUPPLEMENTAL FIGURES

**Supplemental Figure 1. Confirmation of known *rve 4 6 8* phenotypes.** Phenomics experimentation was designed to parallel that resolved by previously by Gray *et al.*, 2017 to ensure growth conditions produced comparable phenotypes prior to molecular experimentation. **(A & B)** Rosette area (mm^2^) and perimeter (mm) over 5 d of growth (15 d – 19 d post-imbibition). Data was extracted using PlantCV (n=20). **(C & D)** Rosette fresh and dry weight (mg) at 28 d post-imbibition (n=10). **(E)** Days to bolting (n=20). Stars denote Student’s t-test *p-value* significance: (***) *p-value* ≤ 0.005 and (*) *p-value* ≤ 0.05.

**Supplemental Figure 2: Diel mRNA gene expression analysis of circadian clock and phytohormone related genes in *rve 4 6 8* and Col-0 plants.** Select genes from different circadian / diel implicated plant cell processes were examined for their expression level at ZT0 and ZT12 by quantitative NanoString expression analysis (see Materials and Methods). **(A)** Expression of circadian clock / RVE8-like related genes *RVE4*, *RVE8*, *PRR7*, *GI* and *CO*. **(B)** Expression of phytohormone genes *GGPPS1*, *ARR7* and *IAA4*. All time-points and genotypes analyzed in biological triplicate (n=3). Stars denote Student’s t-test *p-value* significance: (***) *p-value* ≤ 0.005, (**) *p-value* ≤ 0.01 and (*) *p-value* ≤ 0.05. Primer pairs used in the nanostring analysis are described in Supplemental Data 1.

**Supplemental Figure 3. Association network analysis of significantly changing phosphoproteins maintaining an enriched protein kinase motifs.** The phopshopeptides corresponding to one or more of the 5 enriched phosphorylation motifs were used to generate an association network that relates protein kinase motifs to the underlying protein networks which exhibit genotypic changes in phosphorylation (Supplemental Data 5). Networks were generated using Cytoscape and a combination of the STRING-DB and enhancedGraphics App (see Materials and Methods) using all datatypes and an edge score ≥ 0.7. Protein localization was determined using SUBA4 (https://suba.live/). Highlighted node clusters were manually annotated using a combination of ThaleMine (https://bar.utoronto.ca/thalemine/begin.do) and TAIR (https://www.arabidopsis.org/).

**Supplemental Figure 4:** Additional replicates for the proteasome inhibitor sensitivity growth assay described in Figure 7B.

## SUPPLEMENTAL DATA

**Supplemental Data 1: Primer sets used in Nanostring analysis.**

**Supplemental Data 2: All quantified and significantly changing proteome data.**

**Supplemental Data 3: All quantified and significantly changing phosphoproteome data.**

**Supplemental Data 4: Gene Ontology enrichment analysis.**

**Supplemental Data 5: Comparative analysis of significantly changing phosphorylation sites to significantly cycling phosphorylation sites (JTK-cycle *p-value* ≤ 0.05).**

**Supplemental Data 6: Diel changes in proteasome.**

**Supplemental Data 7: MotifX phosphor-motif enrichment analysis using MoMo MEME suite.**

**Supplemental Data 8: Gas Chromatography Mass Spectrometry (GC-MS) data**

## LITERATURE CITED

Aarabi F, Naake T, Fernie AR, Hoefgen R (2020) Coordinating Sulfur Pools under Sulfate Deprivation. Trends Plant Sci 25: 1227–1239

Abramoff MD, Magalhaes PJ, and Ram SJ (2004) Image Processing with ImageJ. Biophotonics International. 11: 36–42

Akten B, Jauch E, Genova GK, Kim EY, Edery I, Raabe T, Jackson FR (2003) A role for CK2 in the Drosophila circadian oscillator. Nat Neurosci 6: 251–257

Artus NN, Uemura M, Steponkus PL, Gilmour SJ, Lin C, Thomashow MF (1996) Constitutive expression of the cold-regulated Arabidopsis thaliana COR15a gene affects both chloroplast and protoplast freezing tolerance. Proc Natl Acad Sci U S A 93: 13404–9

Astot C, Dolezal K, Nordstrom A, Wang Q, Kunkel T, Moritz T, Chua NH, Sandberg G (2000) An alternative cytokinin biosynthesis pathway. Proc Natl Acad Sci U S A 97: 14778–14783

Baerenfaller K, Massonnet C, Walsh S, Baginsky S, Buhlmann P, Hennig L, Hirsch-Hoffmann M, Howell KA, Kahlau S, Radziejwoski A, Russenberger D, Rutishauser D, Small I, Stekhoven D, Sulpice R, Svozil J, Wuyts N, Stitt M, Hilson P, Granier C, Gruissem W (2012) Systems-based analysis of Arabidopsis leaf growth reveals adaptation to water deficit. Mol Syst Biol 8: 606

Barbier FF, Cao D, Fichtner F, Weiste C, Perez-Garcia MD, Caradeuc M, Le Gourrierec J, Sakr S, Beveridge CA (2021) HEXOKINASE1 signalling promotes shoot branching and interacts with cytokinin and strigolactone pathways. New Phytol 231: 1088–1104

Betti C, Vanhoutte I, Coutuer S, De Rycke R, Mishev K, Vuylsteke M, Aesaert S, Rombaut D, Gallardo R, De Smet F, Xu J, Van Lijsebettens M, Van Breusegem F, Inzé D, Rousseau F, Schymkowitz J, Russinova E (2016) Sequence-Specific Protein Aggregation Generates Defined Protein Knockdowns in Plants. Plant Physiol. 171: 773–87

Bognar LK, Hall A, Adam E, Thain SC, Nagy F, Millar AJ (1999) The circadian clock controls the expression pattern of the circadian input photoreceptor, phytochrome B. Proc Natl Acad Sci U S A 96: 14652–14657

Cervela-Cardona L, Yoshida T, Zhang Y, Okada M, Fernie A, Mas P (2021) Circadian Control of Metabolism by the Clock Component TOC1. Front Plant Sci 12: 683516

Chew YH, Seaton DD, Mengin V, Flis A, Mugford ST, Smith AM, Stitt M, Millar AJ (2017) Linking circadian time to growth rate quantitatively via carbon metabolism. BioRxIv

Chomczynski P, Sacchi N (1987) Single-step method of RNA isolation by acid guanidinium thiocyanate-phenol-chloroform extraction. Anal Biochem 162: 156–159

Chou MF, Schwartz D (2011) Biological sequence motif discovery using motif-x. Curr Protoc Bioinformatics **Chapter 13**: Unit 13 15–24

Choudhary MK, Nomura Y, Shi H, Nakagami H, Somers DE (2016) Circadian profiling of the Arabidopsis proteome using 2D-DIGE. Front. Plant Sci. 7: 1007

Choudhary MK, Nomura Y, Wang L, Nakagami H, Somers DE (2015) Quantitative circadian phosphoproteomic analysis of Arabidopsis reveals extensive clock control of key components in physiological, metabolic and signaling pathways. Mol. Cell. Proteom. 14: 2243–2260

Covington MF, Maloof JN, Straume M, Kay SA, Harmer SL (2008) Global transcriptome analysis reveals circadian regulation of key pathways in plant growth and development. Genome Biol 9: R130

Cox J, Mann M (2008) MaxQuant enables high peptide identification rates, individualized p.p.b.-range mass accuracies and proteome-wide protein quantification. Nat Biotechnol 26: 1367–1372

Creux N, Harmer S (2019) Circadian Rhythms in Plants. Cold Spring Harb Perspect Biol 11

Critchley JH, Zeeman SC, Takaha T, Smith AM, Smith SM (2001) A critical role for disproportionating enzyme in starch breakdown is revealed by a knock-out mutation in Arabidopsis. Plant J. 26: 89–100

Devlin PF, Kay SA (2000) Cryptochromes are required for phytochrome signaling to the circadian clock but not for rhythmicity. Plant Cell 12: 2499–2510

Farinas B, Mas P (2011) Functional implication of the MYB transcription factor RVE8/LCL5 in the circadian control of histone acetylation. Plant J 66: 318–329

Feke A, Liu W, Hong J, Li MW, Lee CM, Zhou EK, Gendron JM (2019) Decoys provide a scalable platform for the identification of plant E3 ubiquitin ligases that regulate circadian function. Elife 8

Feke AM, Hong J, Liu W, Gendron JM (2020) A Decoy Library Uncovers U-Box E3 Ubiquitin Ligases That Regulate Flowering Time in Arabidopsis. Genetics 215: 699–712

Flis A, Mengin V, Ivakov AA, Mugford ST, Hubberten HM, Encke B, Krohn N, Hohne M, Feil R, Hoefgen R, Lunn JE, Millar AJ, Smith AM, Sulpice R, Stitt M (2019) Multiple circadian clock outputs regulate diel turnover of carbon and nitrogen reserves. Plant Cell Environ 42: 549–573

Flis A, Sulpice R, Seaton DD, Ivakov AA, Liput M, Abel C, Millar AJ, Stitt M (2016) Photoperiod-dependent changes in the phase of core clock transcripts and global transcriptional outputs at dawn and dusk in Arabidopsis. Plant Cell Environ 39: 1955–1981

Fukushima A, Kusano M, Nakamichi N, Kobayashi M, Hayashi N, Sakakibara H, Mizuno T, Saito K (2009) Impact of clock-associated Arabidopsis pseudo-response regulators in metabolic coordination. Pr°C Natl Acad Sci U S A 106: 7251–7256

Gehan MA, Fahlgren N, Abbasi A, Berry JC, Callen ST, Chavez L, Doust AN, Feldman MJ, Gilbert KB, Hodge JG, Hoyer JS, Lin A, Liu S, Lizarraga C, Lorence A, Miller M, Platon E, Tessman M, Sax T (2017) PlantCV v2: Image analysis software for high-throughput plant phenotyping. PeerJ 5: e4088

Gibson S, Arondel V, Iba K, Somerville C (1994) Cloning of a temperature-regulated gene encoding a chloroplast omega-3 desaturase from Arabidopsis thaliana. Plant Physiol 106: 1615–1621

Gladman N P, Marshall R S, Lee K-H, Vierstra R D (2016), The Proteasome Stress Regulon Is Controlled by a Pair of NAC Transcription Factors in Arabidopsis, The Plant Cell, Volume 28, Issue 6, Pages 1279–1296, https://doi.org/10.1105/tpc.15.01022

Graf A, Coman D, Uhrig RG, Walsh S, Flis A, Stitt M, Gruissem W (2017) Parallel analysis of Arabidopsis circadian clock mutants reveals different scales of transcriptome and proteome regulation. Open Biol 7

Gray JA, Shalit-Kaneh A, Chu DN, Hsu PY, Harmer SL (2017) The REVEILLE Clock Genes Inhibit Growth of Juvenile and Adult Plants by Control of Cell Size. Plant Physiol 173: 2308–2322

Greenham K, McClung CR (2015) Integrating circadian dynamics with physiological processes in plants. Nat Rev Genet 16: 598–610

Han J-J, Yan X, Wang Q, Tang L, Yu F, Huang X, Wang, Y, Liu J-X, Xie Q (2019), The β5 subunit is essential for intact 26S proteasome assembly to specifically promote plant autotrophic growth under salt stress. New Phytol, 221: 1359–1368. https://doi.org/10.1111/nph.15471

Hardtke CS, Okamoto H, Stoop-Myer C, Deng XW (2002) Biochemical evidence for ubiquitin ligase activity of the Arabidopsis COP1 interacting protein 8 (CIP8). Plant J 30: 385–394

He M, Ding NZ (2020) Plant Unsaturated Fatty Acids: Multiple Roles in Stress Response. Front Plant Sci 11: 562785

Heazlewood JL, Durek P, Hummel J, Selbig J, Weckwerth W, Walther D, Schulze WX (2008) PhosPhAt: a database of phosphorylation sites in Arabidopsis thaliana and a plant-specific phosphorylation site predictor. Nucleic Acids Res 36: D1015–1021

Heazlewood JL, Tonti-Filippini JS, Gout AM, Day DA, Whelan J, Millar AH (2004) Experimental analysis of the Arabidopsis mitochondrial proteome highlights signaling and regulatory components, provides assessment of targeting prediction programs, and indicates plant-specific mitochondrial proteins. Plant Cell 16: 241–256

Hei S, Liu Z, Huang A, She X (2018) The regulator of G-protein signalling protein mediates D-glucose-induced stomatal closure via triggering hydrogen peroxide and nitric oxide production in Arabidopsis. Funct Plant Biol 45: 509–518

Herold A, Lewis DH (1977) MANNOSE AND GREEN PLANTS: OCCURRENCE, PHYSIOLOGY AND METABOLISM, AND USE AS A TOOL TO STUDY THE ROLE OF ORTHOPHOSPHATE. New Phytologist 79: 1–40

Hill CB, Roessner U (2013) Metabolic Profiling of Plants by GC–MS. The Handbook of Plant Metabolomics

Hsu PY, Devisetty UK, Harmer SL (2013) Accurate timekeeping is controlled by a cycling activator in Arabidopsis. Elife 2: e00473

Huang S, Li L, Petereit J, Millar AH (2020) Protein turnover rates in plant mitochondria. Mitochondrion 53: 57–65

Huang X, Zhang X, Gong Z, Yang S, Shi Y (2017) ABI4 represses the expression of type-A ARRs to inhibit seed germination in Arabidopsis. Plant J 89: 354–365

Jaglo-Ottosen KR, Gilmour SJ, Zarka DG, Schabenberger O, Thomashow MF (1998) Arabidopsis CBF1 overexpression induces COR genes and enhances freezing tolerance. Science 280: 104–6.

Kidokoro S, Hayashi K, Haraguchi H, Ishikawa T, Soma F, Konoura I, Toda S, Mizoi J, Suzuki T, Shinozaki K, Yamaguchi-Shinozaki K (2021) Posttranslational regulation of multiple clock-related transcription factors triggers cold-inducible gene expression in Arabidopsis. Proc Natl Acad Sci U S A 118: e2021048118

Kotchoni SO, Larrimore KE, Mukherjee M, Kempinski CF, Barth C (2009) Alterations in the endogenous ascorbic acid content affect flowering time in Arabidopsis. Plant Physiol 149: 803–815

Krahmer J, Hindle M, Perby LK, Nielson TH, VanOoijen G, Halliday KJ, Le Bihan T, Millar AJ (2022) Circadian protein regulation in the green lineage II. The clock gene circuit controls a phospho-dawn in Arabidopsis thaliana. Mol. Cell. Proteom 21: 100172

Kurepa J, Wang S, Li Y, Zaitlin D, Pierce JA, Smalle JA (2009) Loss of 26S Proteasome Function Leads to Increased Cell Size and Decreased Cell Number in Arabidopsis Shoot Organs. Plant Physiology 150: 178–189

Kurepa J, Wang S, Li Y, Smalle JA (2009) Proteasome regulation, plant growth and stress tolerance. Plant Signalling and Behaviour 4: 924–927

Lee SH, Li CW, Koh KW, Chuang HY, Chen YR, Lin CS, Chan MT (2014) MSRB7 reverses oxidation of GSTF2/3 to confer tolerance of Arabidopsis thaliana to oxidative stress. J Exp Bot 65: 5049–5062

Leschevin M, Ismael M, Quero A, San Clemente H, Roulard R, Bassard S, Marcelo P, Pageau K, Jamet E, Rayon C (2021) Physiological and Biochemical Traits of Two Major Arabidopsis Accessions, Col-0 and Ws, Under Salinity. Front Plant Sci 12: 639154

Leutert M, Rodriguez-Mias RA, Fukuda NK, Villen J (2019) R2-P2 rapid-robotic phosphoproteomics enables multidimensional cell signaling studies. Mol Syst Biol 15: e9021

Li L, Nelson CJ, Trosch J, Castleden I, Huang S, Millar AH (2017) Protein Degradation Rate in Arabidopsis thaliana Leaf Growth and Development. Plant Cell 29: 207–228

Liu XS, Yang X, Wang LM, Duan QQ, Huang DF (2016) Comparative analysis of metabolites profile in spinach (Spinacia oleracea L.) affected by different concentrations of gly and nitrate. Scientia Horticulturae 204: 8–15

Lu H, McClung CR, Zhang C (2017) Tick Tock: Circadian Regulation of Plant Innate Immunity. Annu Rev Phytopathol 55: 287–311

Lu SX, Liu H, Knowles SM, Li J, Ma L, Tobin EM, Lin C (2011) A role for protein kinase casein kinase2 alpha-subunits in the Arabidopsis circadian clock. Plant Physiol 157: 1537–1545

Marti Ruiz MC, Jung HJ, Webb AAR (2020) Circadian gating of dark-induced increases in chloroplast- and cytosolic-free calcium in Arabidopsis. New Phytol 225: 1993–2005

Maszkowska, J, Dębski, J, Kulik, A, et al. (2019) Phosphoproteomic analysis reveals that dehydrins ERD10 and ERD14 are phosphorylated by SNF1-related protein kinase 2.10 in response to osmotic stress. Plant Cell Environ. 42: 931–946

McClung CR (2019) The Plant Circadian Oscillator. Biology (Basel) 8: 14

Mehlmer N, Wurzinger B, Stael S, Hofmann-Rodrigues D, Csaszar E, Pfister B, Bayer R, Teige M (2010) The Ca(2+) -dependent protein kinase CPK3 is required for MAPK-independent salt-stress acclimation in Arabidopsis. Plant J 63: 484–498

Mehta D, Ghahremani M, Perez-Fernandez M, Tan M, Schlapfer P, Plaxton WC, Uhrig RG (2021) Phosphate and phosphite have a differential impact on the proteome and phosphoproteome of Arabidopsis suspension cell cultures. Plant J 105: 924–941

Mehta D, Krahmer J, Uhrig RG (2021) Closing the protein gap in plant chronobiology. Plant J 106: 1509–1522

Mehta D, Scandola S, Uhrig RG (2022) BoxCar and Library-Free Data-Independent Acquisition Substantially Improve the Depth, Range, and Completeness of Label-Free Quantitative Proteomics. Anal Chem 94: 793–802

Mori IC, Murata Y, Yang Y, Munemasa S, Wang YF, Andreoli S, Tiriac H, Alonso JM, Harper JF, Ecker JR, Kwak JM, Schroeder JI (2006) CDPKs CPK6 and CPK3 function in ABA regulation of guard cell S-type anion- and Ca(2+)-permeable channels and stomatal closure. PLoS Biol 4: e327

Mulekar JJ, Huq E (2014) Expanding roles of protein kinase CK2 in regulating plant growth and development. J Exp Bot 65: 2883–2893

Nakamichi N (2020) The Transcriptional Network in the Arabidopsis Circadian Clock System. Genes (Basel) 11

Nohales MA, Kay SA (2016) Molecular mechanisms at the core of the plant circadian oscillator. Nat Struct Mol Biol 23: 1061–1069

Patterson JA, Tetlow IJ, Emes MJ (2018) Bioinformatic and in vitro Analyses of Arabidopsis Starch Synthase 2 Reveal Post-translational Regulatory Mechanisms. Front Plant Sci 9: 1338

Purzner T, Purzner J, Buckstaff T, Cozza G, Gholamin S, Rusert JM, Hartl TA, Sanders J, Conley N, Ge X, Langan M, Ramaswamy V, Ellis L, Litzenburger U, Bolin S, Theruvath J, Nitta R, Qi L, Li XN, Li G, Taylor MD, Wechsler-Reya RJ, Pinna LA, Cho YJ, Fuller MT, Elias JE, Scott MP (2018) Developmental phosphoproteomics identifies the kinase CK2 as a driver of Hedgehog signaling and a therapeutic target in medulloblastoma. Sci Signal 11: eaau5147

Qi T, Liu Z, Fan M, Chen Y, Tian H, Wu D, Gao H, Ren C, Song S, Xie D (2017) GDP-D-mannose epimerase regulates male gametophyte development, plant growth and leaf senescence in Arabidopsis. Sci Rep 7: 10309

Qin N, Xu D, Li J, Deng XW (2020) COP9 signalosome: Discovery, conservation, activity and, function. JIPB 62: 90–103

Rawat R, Takahashi N, Hsu PY, Jones MA, Schwartz J, Salemi MR, Phinney BS, Harmer SL (2011) REVEILLE8 and PSEUDO-REPONSE REGULATOR5 form a negative feedback loop within the Arabidopsis circadian clock. PLoS Genet 7: e1001350

Rodriguez MC, Mehta D, Tan M, Uhrig RG (2021) Quantitative Proteome and PTMome Analysis of Arabidopsis thaliana Root Responses to Persistent Osmotic and Salinity Stress. Plant Cell Physiol 62: 1012–1029

Ruiz-Sola MA, Barja MV, Manzano D, Llorente B, Schipper B, Beekwilder J, Rodriguez-Concepcion M (2016) A Single Arabidopsis Gene Encodes Two Differentially Targeted Geranylgeranyl Diphosphate Synthase Isoforms. Plant Physiol 172: 1393–1402

Sadandom A, Bailey M, Ewan R, Lee J, Neils S (2012) The ubiquitin-proteasome system: central modifier of plant signalling. New Phytologist 196: 13–28.

Sanchez SE, Rugnone ML, Kay SA (2020) Light Perception: A Matter of Time. Mol Plant 13: 363–385

Senkler J, Senkler M, Eubel H, Hildebrandt T, Lengwenus C, Schertl P, Schwarzlander M, Wagner S, Wittig I, Braun HP (2017) The mitochondrial complexome of Arabidopsis thaliana. Plant J 89: 1079–1092

Seok HY, Nguyen LV, Van Nguyen D, Lee SY, Moon YH (2020) Investigation of a Novel Salt Stress-Responsive Pathway Mediated by Arabidopsis DEAD-Box RNA Helicase Gene AtRH17 Using RNA-Seq Analysis. Int J Mol Sci 21: 1595

Sharma V, Ichikawa M, Freeze HH (2014) Mannose metabolism: more than meets the eye. Biochem Biophys Res Commun 453: 220–228

Shim JS, Kubota A, Imaizumi T (2017) Circadian Clock and Photoperiodic Flowering in Arabidopsis: CONSTANS Is a Hub for Signal Integration. Plant Physiol 173: 5–15

Simon NML, Graham CA, Comben NE, Hetherington AM, Dodd AN (2020) The Circadian Clock Influences the Long-Term Water Use Efficiency of Arabidopsis. Plant Physiol 183: 317–330

Smalle J, Kurepa J, Yang P, Babiychuk E, Kushnir S, Durski A, Vierstra R D (2002) Cytokinin Growth Responses in Arabidopsis Involve the 26S Proteasome Subunit RPN12. The Plant Cell, 14 1: 17–32

Smith AM, Zeeman SC (2006) Quantification of starch in plant tissues. Nature Protocols 1: 1342–1345

Smith AM, Zeeman SC (2020) Starch: A Flexible, Adaptable Carbon Store Coupled to Plant Growth. Annu Rev Plant Biol 71: 217–245

Somers DE, Devlin PF, Kay SA (1998) Phytochromes and cryptochromes in the entrainment of the Arabidopsis circadian clock. Science 282: 1488–1490

Streb S, Zeeman SC (2012) Starch metabolism in Arabidopsis. Arabidopsis Book 10: e0160

Sugano S, Andronis C, Green RM, Wang ZY, Tobin EM (1998) Protein kinase CK2 interacts with and phosphorylates the Arabidopsis circadian clock-associated 1 protein. Proc Natl Acad Sci U S A 95: 11020–11025

Tamaru T, Hirayama J, Isojima Y, Nagai K, Norioka S, Takamatsu K, Sassone-Corsi P (2009) CK2alpha phosphorylates BMAL1 to regulate the mammalian clock. Nat Struct Mol Biol 16: 446–448

Tetlow IJ, Wait R, Lu Z, Akkasaeng R, Bowsher CG, Esposito S, Kosar-Hashemi B, Morell MK, Emes MJ (2004) Protein phosphorylation in amyloplasts regulates starch branching enzyme activity and protein-protein interactions. Plant Cell 16: 694–708

Thalmann M, Pazmino D, Seung D, Horrer D, Nigro A, Meier T, Kolling K, Pfeifhofer HW, Zeeman SC, Santelia D (2016) Regulation of Leaf Starch Degradation by Abscisic Acid Is Important for Osmotic Stress Tolerance in Plants. Plant Cell 28: 1860–1878

Toledo M, Batista-Gonzalez A, Merheb E, Aoun ML, Tarabra E, Feng D, Sarparanta J, Merlo P, Botre F, Schwartz GJ, Pessin JE, Singh R (2018) Autophagy Regulates the Liver Clock and Glucose Metabolism by Degrading CRY1. Cell Metab 28: 268–281 e264

Torres-Franklin ML, Repellin A, Huynh VB, d’Arcy-Lameta A, Zuily-Fodil Y, Thi ATP (2009) Omega-3 fatty acid desaturase (FAD3, FAD7, FAD8) gene expression and linolenic acid content in cowpea leaves submitted to drought and after rehydration. Environmental and Experimental Botany 65: 162–169

Tyanova S, Temu T, Sinitcyn P, Carlson A, Hein MY, Geiger T, Mann M, Cox J (2016) The Perseus computational platform for comprehensive analysis of (prote)omics data. Nat Methods 13: 731–740

Uehara TN, Mizutani Y, Kuwata K, Hirota T, Sato A, Mizoi J, Takao S, Matsuo H, Suzuki T, Ito S, Saito AN, Nishiwaki-Ohkawa T, Yamaguchi-Shinozaki K, Yoshimura T, Kay SA, Itami K, Kinoshita T, Yamaguchi J, Nakamichi N (2019) Casein kinase 1 family regulates PRR5 and TOC1 in the Arabidopsis circadian clock. Proc Natl Acad Sci U S A 116: 11528–11536

Uhrig RG, Echevarria-Zomeno S, Schlapfer P, Grossmann J, Roschitzki B, Koerber N, Fiorani F, Gruissem W (2021) Diurnal dynamics of the Arabidopsis rosette proteome and phosphoproteome. Plant Cell Environ 44: 821–841

Uhrig RG, Schlapfer P, Roschitzki B, Hirsch-Hoffmann M, Gruissem W (2019) Diurnal changes in concerted plant protein phosphorylation and acetylation in Arabidopsis organs and seedlings. Plant J 99: 176–194

Upchurch RG (2008) Fatty acid unsaturation, mobilization, and regulation in the response of plants to stress. Biotechnol Lett 30: 967–977

Usmani MM, Nawaz F, Majeed S, Shehzad MA, Ahmad KS, Akhtar G, Aqib M, Shabbir RN (2020) Sulfate-mediated Drought Tolerance in Maize Involves Regulation at Physiological and Biochemical Levels. Sci Rep 10: 1147

Üstün, S. and Börnke, F. (2017). Ubiquitin Proteasome Activity Measurement in Total Plant Extracts. Bio-protocol 7(17): e2532.

Veremeichik GN, Shkryl YN, Gorpenchenko TY, Silantieva SA, Avramenko TV, Brodovskaya EV, Bulgakov VP (2021) Inactivation of the auto-inhibitory domain in Arabidopsis AtCPK1 leads to increased salt, cold and heat tolerance in the AtCPK1-transformed Rubia cordifolia L cell cultures. Plant Physiol Biochem 159: 372–382

Wang W, Paik I, Kim J, Hou X, Sung S, Huq E (2021) Direct phosphorylation of HY5 by SPA kinases to regulate photomorphogenesis in Arabidopsis. New Phytol 230: 2311–2326

Wasternack C, Hause B (2019) The missing link in jasmonic acid biosynthesis. Nat Plants 5: 776–777

Zhang M, Barg R, Yin M, Gueta-Dahan Y, Leikin-Frenkel A, Salts Y, Shabtai S, Ben-Hayyim G (2005) Modulated fatty acid desaturation via overexpression of two distinct omega-3 desaturases differentially alters tolerance to various abiotic stresses in transgenic tobacco cells and plants. Plant J 44: 361–371

Zhu W, Zhou H, Lin F, Zhao X, Jiang Y, Xu D, Deng XW (2020) COLD-REGULATED GENE27 Integrates Signals from Light and the Circadian Clock to Promote Hypocotyl Growth in Arabidopsis. Plant Cell 32: 3155–3169

Zhu, Y., Huang, P., Guo, P., Chong, L., Yu, G., Sun, X., Hu, T., Li, Y., Hsu, C.-C., Tang, K., Zhou, Y., Zhao, C., Gao, W., Tao, W.A., Mengiste, T. and Zhu, J.-K. (2020), CDK8 is associated with RAP2.6 and SnRK2.6 and positively modulates abscisic acid signaling and drought response in Arabidopsis. New Phytol, 228: 1573–1590.

