## Supplementary figures and images for "Multi-omic analysis of the Arabidopsis clock activator mutant *rve 4 6 8* reveals connections to carbohydrate metabolism and proteasome regulation"

### Supplemental Figure 1

#### Supplemental Figure 1

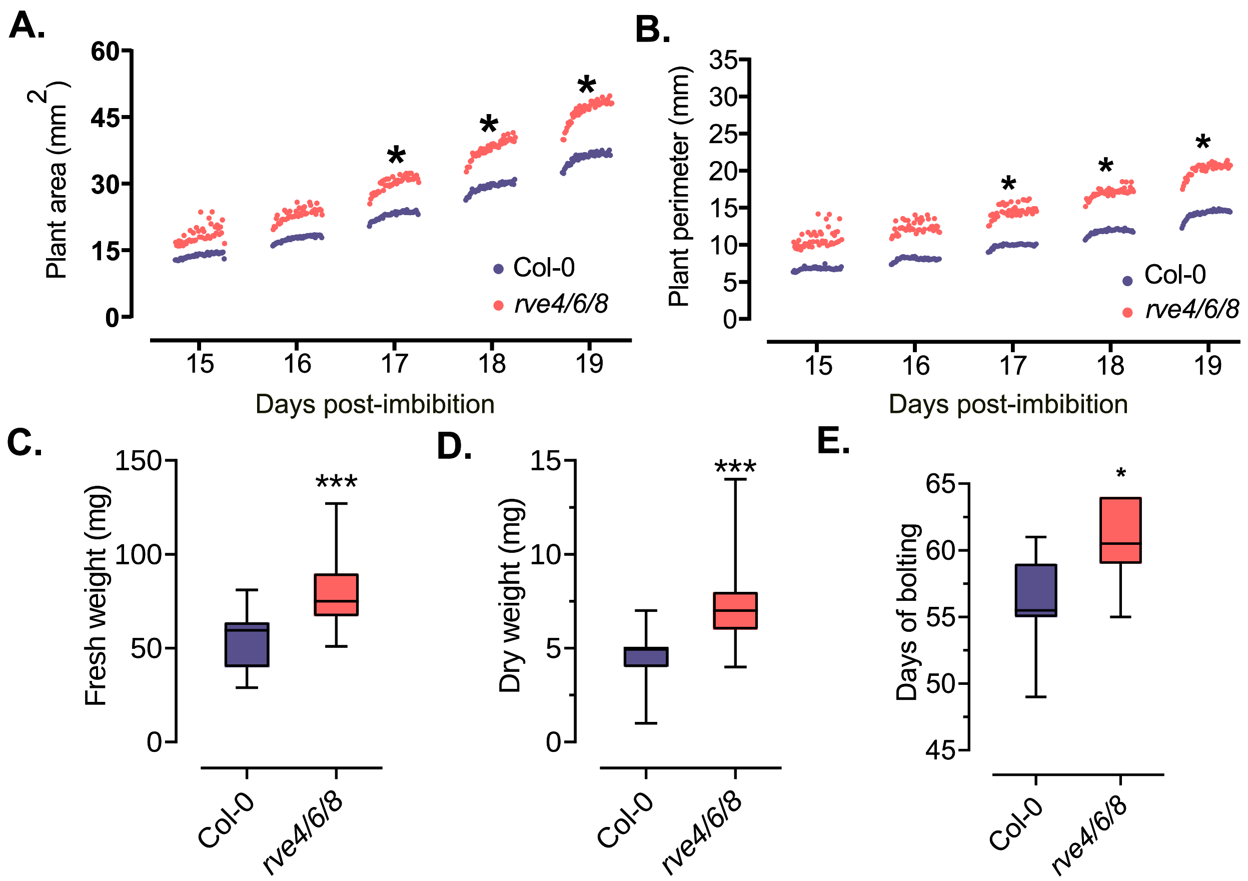

### Supplemental Figure 2

#### Supplemental Figure 2

##### **A.** Circadian Clock / RVE8-Like Related

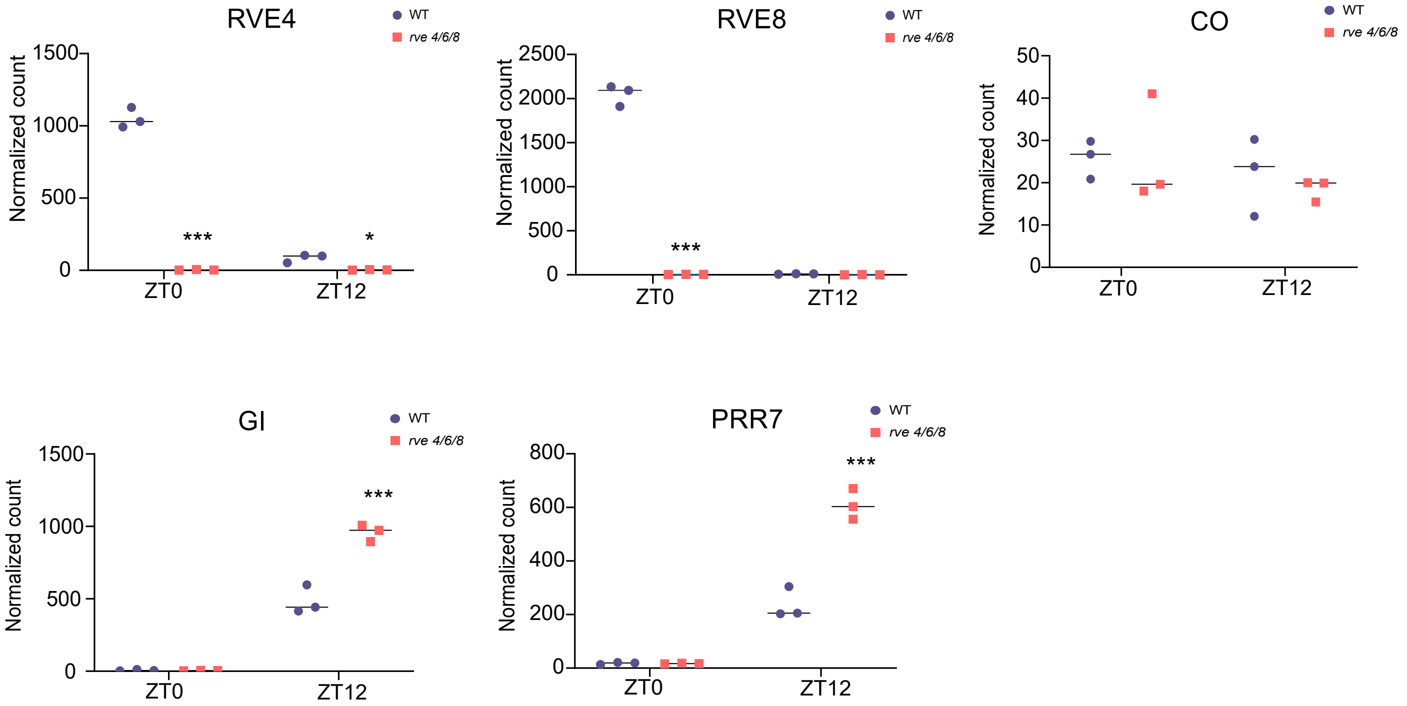


##### **B.** Isoprenoid/ Cytokinin/ Auxin Related

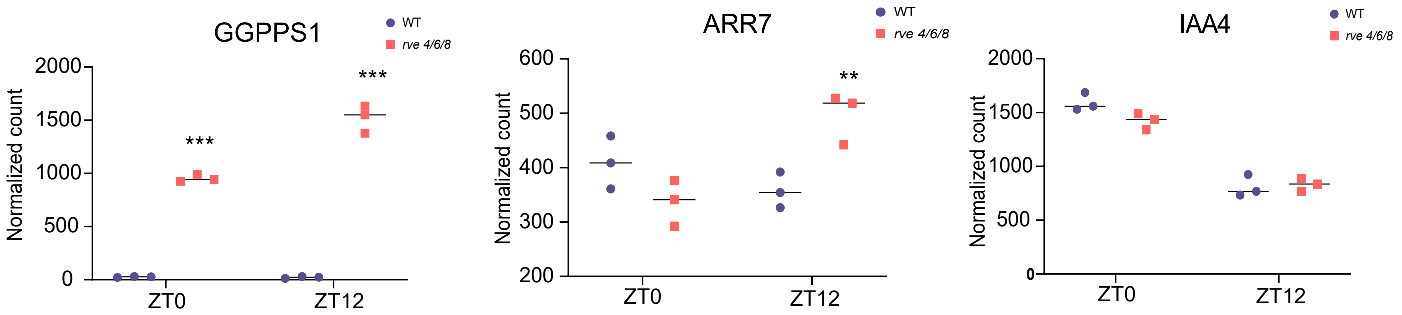

### Supplemental Figure 3

#### Supplemental Figure 3

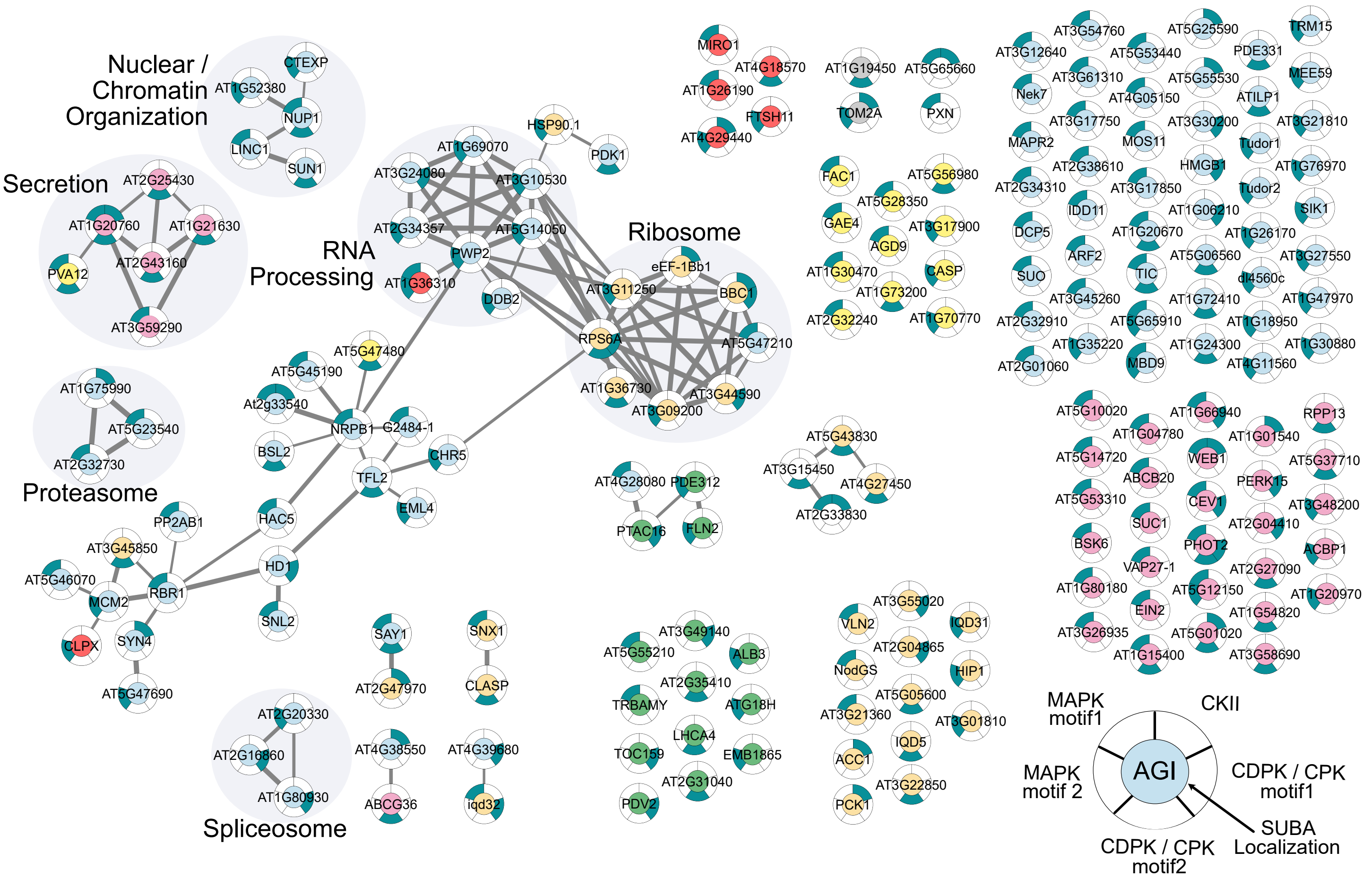

### Supplemental Figure 4

**Supplemental Figure 4**

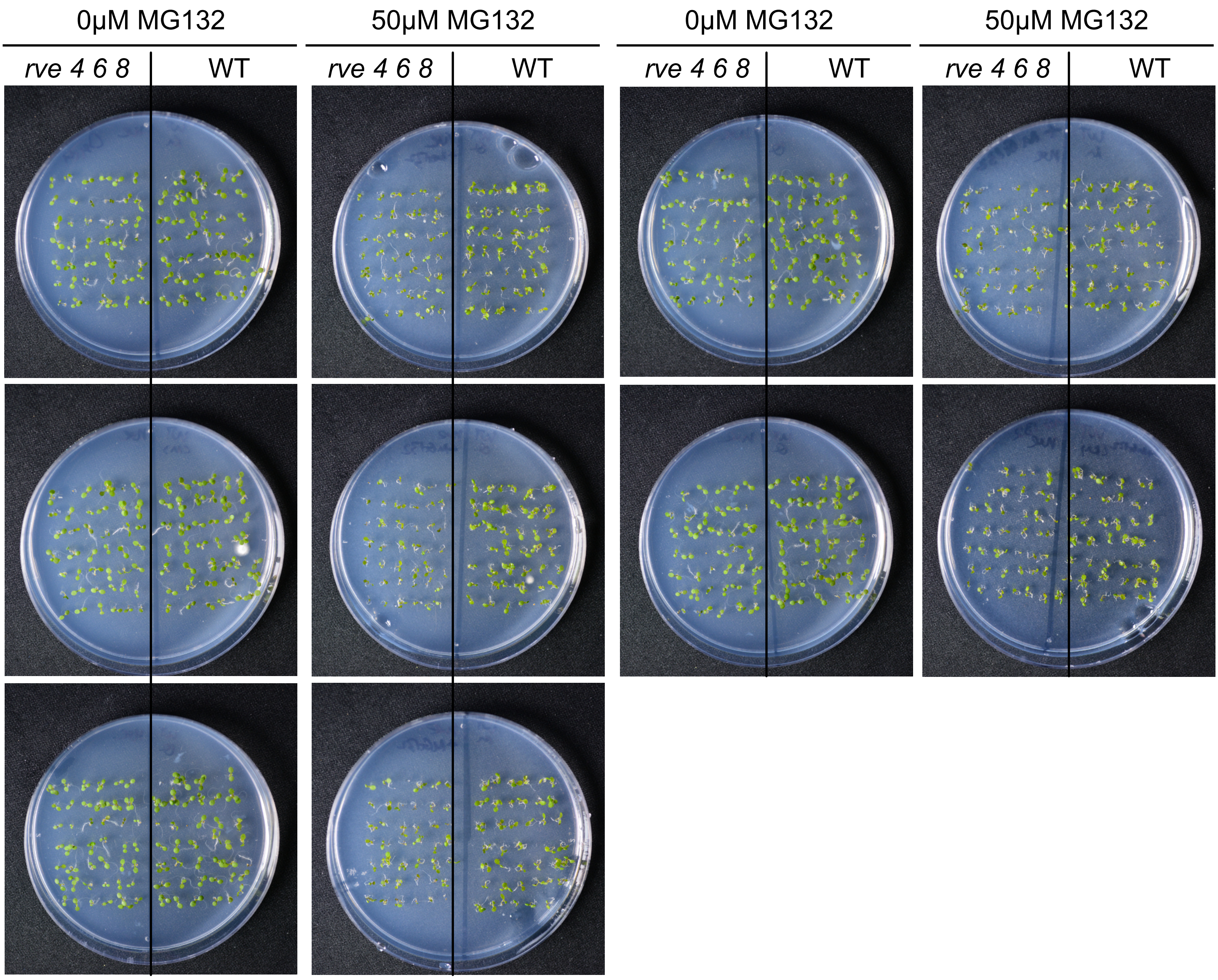
